# Release of Histone H3K4-reading transcription factors from chromosomes in mitosis is independent of adjacent H3 phosphorylation

**DOI:** 10.1101/2023.02.28.530230

**Authors:** Rebecca J. Harris, Maninder Heer, Mark D. Levasseur, Tyrell N. Cartwright, Bethany Weston, Jennifer L. Mitchell, Jonathan M. Coxhead, Luke Gaughan, Lisa Prendergast, Daniel Rico, Jonathan M.G. Higgins

## Abstract

Histone modifications influence the recruitment of reader proteins to chromosomes to regulate events including transcription and cell division. The idea of a histone code, where particular combinations of modifications specify unique downstream functions, is widely accepted and can be demonstrated *in vitro*. For example, on synthetic peptides, phosphorylation of Histone H3 at threonine-3 (H3T3ph) prevents the binding of reader proteins that recognise trimethylation of the adjacent lysine-4 (H3K4me3), including the TAF3 component of TFIID. To study these combinatorial effects in cells, we analyzed the genome-wide distribution of H3T3ph and H3K4me3 during mitosis. We find that H3K4me3 hinders adjacent H3T3ph deposition in cells, and that the PHD domain of TAF3 can bind H3K4me3 in mitotic chromatin despite the presence of H3T3ph. Unlike *in vitro*, H3K4 readers are displaced from chromosomes in mitosis in Haspin-depleted cells lacking H3T3ph. H3T3ph is therefore unlikely to be responsible for transcriptional downregulation during cell division.

## INTRODUCTION

Chromatin undergoes dramatic changes to facilitate cell division. Interphase chromosomes are organized in loops, topologically associating domains (TADs), and compartments that allow cell-type-specific gene expression programmes to be carried out (Gibcus and Dekker, 2013). However, these structures are rapidly lost as cells enter mitosis and the chromosomes condense (Gibcus et al., 2018; Ginno et al., 2018; Nagano et al., 2017; Naumova et al., 2013; Oomen et al., 2019; Pelham-Webb et al., 2021; Raccaud et al., 2019; Teves et al., 2016; Zhang et al., 2019). While many proteins are recruited to chromatin to regulate chromosome condensation, cohesion, and segregation, others are displaced during mitosis. For example, numerous transcription factors lose sequence-specific associations with chromosomes and transcription is minimized, and the recruitment of DNA damage response proteins is curtailed (Blackford and Stucki, 2020; Djeghloul et al., 2020; Fonseca et al., 2012; Ginno et al., 2018; Gottesfeld and Forbes, 1997; Konrad, 1963; Liang et al., 2015; Martinez-Balbas et al., 1995; Oomen et al., 2019; Palozola et al., 2017; Raccaud et al., 2019; Shermoen and O’Farrell, 1991; Taylor, 1960; Teves et al., 2016). As cells re-enter G1, interphase structures progressively reform and transcription factors must re-associate to re-initiate gene expression. These findings raise a number of important questions, including how the association of transcription factors with chromatin is regulated during the cell cycle, and how appropriate gene expression patterns are re-established after cell division. Knowledge of how these changes are controlled is vital to understand how accurate chromosome segregation takes place and how interphase functions are properly restored.

In one view, the transmission of transcription factors from mother to daughter cells, even if they are dissociated from chromatin, might be sufficient to transfer memory of the gene transcription programme (Ptashne, 2007). Nevertheless, it is widely believed that chromatin-based mechanisms provide a means to regulate gene expression during cell division. In this scenario, although many transcription factors may be displaced, “bookmarks” carried on chromosomes in mitosis contribute to the inheritance of gene expression patterns, or to the speed at which gene transcription is re-initiated. Indeed, the increased DNA accessibility of active promoters is largely maintained in mitosis (Festuccia et al., 2019; Gazit et al., 1982; Ginno et al., 2018; Hsiung et al., 2015; Martinez-Balbas et al., 1995; Michelotti et al., 1997; Oomen et al., 2019; Pelham-Webb et al., 2021; Teves et al., 2016). Various bookmarks have been proposed, including the retention of transcription factors at a subset of specific binding sites, continued basal levels of transcription, and post translational modifications of DNA and histones (Ito and Zaret, 2022; Wang and Higgins, 2013).

Histone modifications constitute an appealing system to provide a bookmarking mechanism. Many such modifications are implicated in the regulation of gene expression during interphase. While these major regulatory modifications are often recognized and dynamically bound by reader proteins, the modifications themselves are often more stable, with some persisting throughout the cell cycle (Behera et al., 2019; Javasky et al., 2018; Kang et al., 2020; Kouskouti and Talianidis, 2005; Liu et al., 2017; Pelham-Webb et al., 2021; Valls et al., 2005). One example is H3K4me3, a mark characteristic of active promoters (Javasky et al., 2018; Kang et al., 2020; Kelly et al., 2010b; Kouskouti and Talianidis, 2005; Liang et al., 2015; Liu et al., 2017; Oomen et al., 2019; Valls et al., 2005). Other histone modifications are relatively dynamic and some, including numerous phosphorylation marks, are strongly upregulated during mitosis. Examples include phosphorylation of H3T3, H3S10, H3S28 and H2AT120 (Dai et al., 2005; Wang and Higgins, 2013). One function of these phosphorylation marks is the recruitment of proteins to mitotic chromosomes. For instance, H3T3ph and H2AT120ph bring the Chromosomal Passenger Complex (CPC) to centromeres (Kawashima et al., 2010; Kelly et al., 2010a; Wang et al., 2010; Yamagishi et al., 2010).

Histone modifications that are retained from interphase through mitosis could allow local chromatin states to be transmitted through cell division (Behera et al., 2019; Hsiung et al., 2016; Javasky et al., 2018; Kang et al., 2020; Kouskouti and Talianidis, 2005; Liang et al., 2015; Liu et al., 2017; Oomen et al., 2019; Pelham-Webb et al., 2021; Valls et al., 2005). Whether reader proteins that recognise these histone marks are also required for bookmarking is less clear. For example, the acetylated-histone reading protein BRD4 is retained on chromatin in mitosis and was originally proposed to be involved in bookmarking, but more recent work suggests that it is the underlying histone marks that are critical (Behera et al., 2019; Dey et al., 2009; Zhao et al., 2011). Consistent with this, numerous proteins that read the histone methylation status at H3K4, H3K9, and H3K27 appear to be substantially displaced from chromosomes in mitosis (Djeghloul et al., 2020; Fischle et al., 2005; Fonseca et al., 2012; Gatchalian et al., 2013; Gatchalian et al., 2016; Hirota et al., 2005). Among these proteins is TAF3, a subunit of the general transcription factor complex TFIID, which recognizes H3K4me2/3 (characteristic of active promoters) through its PHD finger (Lauberth et al., 2013; van Nuland et al., 2013; Varier et al., 2010; Vermeulen et al., 2007). TFIID is a component of the RNA polymerase II preinitiation complex and essential for transcription. Because of this, the displacement of TFIID from H3K4me2/3-marked histones during mitosis has been associated with the general downregulation of transcription during cell division (Gatchalian et al., 2016; Varier et al., 2010).

The striking observation that many major regulatory histone modification sites (e.g. H3K4, H3K9, H3K27 and H2AK119) are adjacent to sites that can be phosphorylated in mitosis led Fischle *et al*. to propose the existence of methyl-phos switches (Fischle et al., 2003). In this model, phosphorylation would prevent the proteins that bind to the adjacent methylated lysine from associating with chromatin.

Indeed, subsequent work suggested that H3S10 phosphorylation by Aurora B displaces the H3K9me3-binding Heterochromatin Protein 1 (HP1) from chromatin in mitosis (Fischle et al., 2005; Hirota et al., 2005). It was proposed that methyl-phos switches may be a widespread mechanism to regulate chromatin association of histone reading proteins.

A significant example is H3K4me3 and the adjacent H3T3, a residue which is phosphorylated during mitosis by the kinase Haspin (Dai et al., 2005). In the methyl-phos switch model, H3T3 phosphorylation in mitosis would displace reader proteins that bind H3K4me3 at promoters, thus contributing to transcriptional repression, while preserving a memory of gene activation (because the H3K4me3 mark persists throughout mitosis). In line with this, *in vitro* experiments show that more than thirty H3K4 reader proteins, including TAF3 (Gatchalian et al., 2016; Kungulovski et al., 2016; Shanle et al., 2017; Varier et al., 2010), can no longer bind to a peptide that also harbours H3T3ph (see Table S1). Methyl-phos switching was therefore proposed to be a major mechanism to regulate transcription during cell division and could modulate protein dissociation from chromatin in a locus-specific manner.

This model, however, is based largely on *in vitro* observations. Notably, methyl-phos switching requires that methylation and phosphorylation co-occur *in vivo*, but little is known about the precise genome-wide localization of histone phosphorylation in mitosis. Previous studies provided some evidence for H3T3/K4 switching in cells (Ali et al., 2013; Gatchalian et al., 2013; Gatchalian et al., 2016; Noh et al., 2015; Quadri et al., 2021; Varier et al., 2010), but often rely on the inverse correlation of H3T3ph and H3K4 reader protein levels on chromatin in microscopy experiments. Here, we analyze the genome-wide distribution and function of H3T3ph and find that the release of H3K4-reading transcription factors from chromosomes in mitosis is independent of methyl-phos switching.

## RESULTS

### Determining the specificity of antibodies recognizing H3T3ph and H3K4me2/3

We wished to determine the localization of H3T3ph genome-wide in mitosis using chromatin immunoprecipitation and next generation sequencing (ChIP-seq), and to compare it to H3K4me3. As H3T3 and H3K4 are adjacent residues in Histone H3, it was critical to use antibodies specific for the histone marks in question that are not blocked by each other or by additional modifications in nearby amino acids. Therefore, we characterised the specificity of H3T3ph and H3K4me3 antibodies before performing ChIP-seq experiments.

Previously, we have found that the affinity-purified rabbit polyclonal antibody B8634 selectively binds H3T3ph *in vitro* and in cells, and that RNAi, inhibition, or CRISPR/Cas9 knockout of the H3T3 kinase Haspin eliminates epitope recognition in cells (Cartwright et al., 2022; Dai et al., 2005; Wang et al., 2012). Using ELISAs on chemically synthesized Histone H3 peptides, we found that the B8634 antibody was indeed specific for H3T3ph, and did not recognize H3S10ph, H3T11ph, or H3T22ph (Figure S1A).

Importantly, peptides carrying H3K4me1, K4me2, or K4me3 in addition to H3T3ph were recognized similarly to peptides carrying H3T3ph alone, showing that adjacent H3K4 methylation did not block recognition of H3T3ph (Figure 1A).

**Figure 1.**
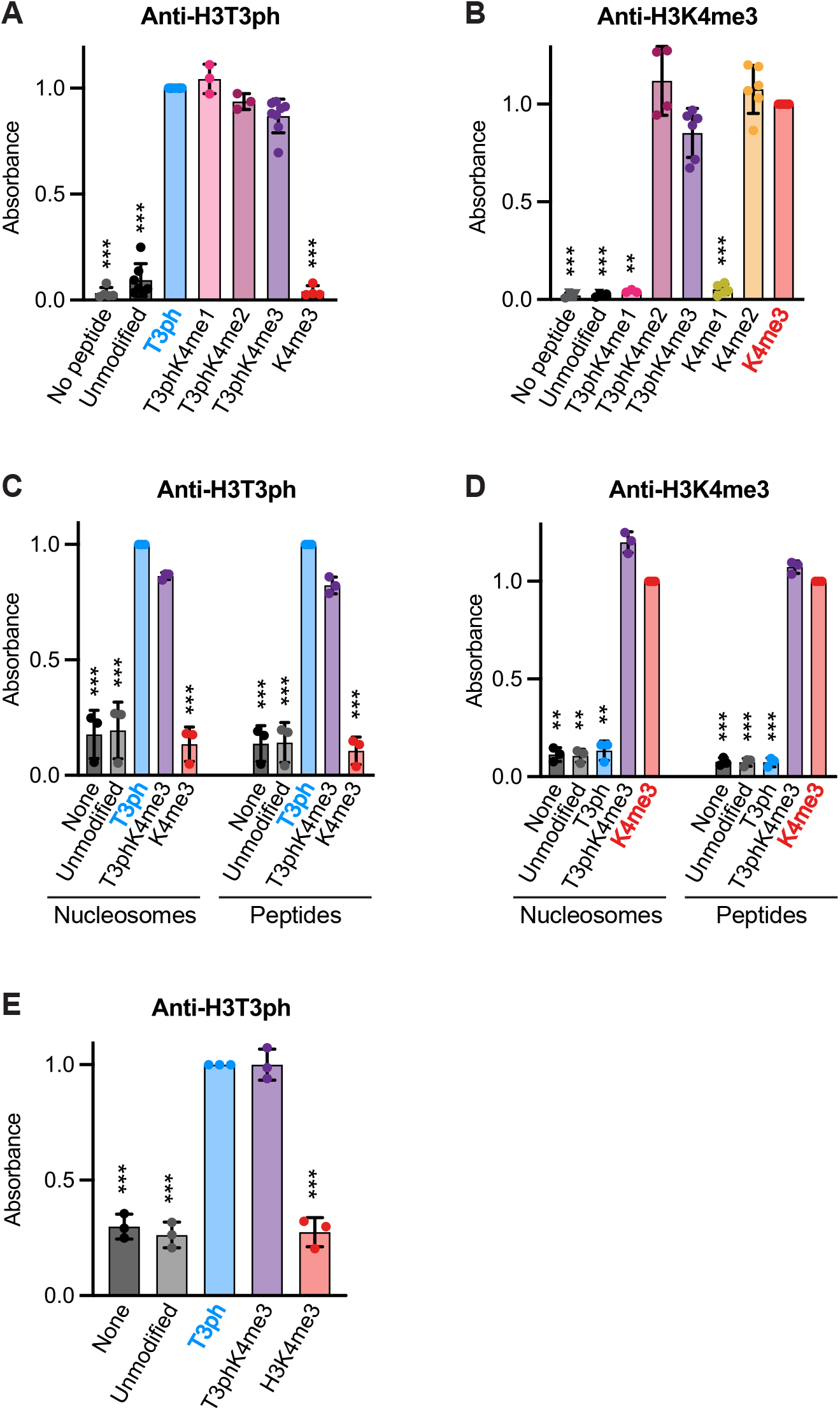
Antibody characterisation by peptide and nucleosome ELISA. **A**. H3T3ph antibody B8634 binding to various H3 peptides (n = 3 to 8). **B**. H3K4me3 antibody C42D8 binding to various H3 peptides (n = 3 to 6). **C**. H3T3ph antibody B8634 binding to peptide and nucleosomal substrates (n = 3). **D**. H3K4me3 antibody C42D8 binding to peptide and nucleosomal substrates (n = 3). **E**. H3T3ph antibody B8634 binding to nucleosomes in the presence of sheared chromatin (n = 3). Data were normalized to the mean signal of the antibodies binding the expected target peptide or nucleosome (i.e. H3T3ph or H3K4me3). Bars represent mean ± SD. Statistical analysis was carried out using non-normalised data, *** p < 0.0001, ** p < 0.001, * p < 0.01, when compared to binding to the expected modification (H3T3ph or H3K4me3). See also Figure S1.

Likewise, the rabbit monoclonal H3K4me3 antibody C42D8 did not recognize H3K9, H3K14, H3K23 or H3K27 trimethylation (Figure S1B), but it did bind to both H3K4me2 and H3K4me3 peptides, and recognition of H3K4me2/3 was not significantly altered by adjacent H3T3ph (Figure 1B) Extensive assessment of C42D8 by others confirms these results (Rothbart et al., 2015; Shah et al., 2018).

Because there is evidence that testing antibodies with histone peptides does not fully recapitulate their recognition properties in ChIP-seq experiments (Shah et al., 2018), we tested them using recombinant nucleosomes containing H3T3ph, H3K4me3, or both marks together. Again, the antibodies showed specificity for the expected single modifications, and both similarly recognized the H3T3phK4me3 dually-modified nucleosomes (Figure 1C, D). We also confirmed that the H3T3ph antibody recognized H3T3ph and H3T3phK4me3-containing nucleosomes equally in the presence of sheared mitotic chromatin, conditions closely approximating those of a ChIP-seq experiment (Figure 1E).

Finally, we used immunoblotting to confirm that the B8634 H3T3ph and C42D8 H3K4me2/3 antibodies recognized predominantly an H3-sized protein of approximately 17 kDa in HeLa cell lysates, and that H3T3ph was detected in mitotic but not asynchronously growing cells, whereas H3K4me2/3 was found at similar levels in both cases (Figure S1C). From these results we conclude that the B8634 H3T3ph antibody recognizes H3T3ph specifically whether or not H3K4 is methylated, and that the C42D8 antibody is suitable to monitor H3K4me2/3 regardless of the presence of H3T3ph.

### Genome-wide H3T3ph and H3K4me2/3 distribution in interphase and mitosis

To determine the distribution of H3T3ph and H3K4me2/3 in mitosis using ChIP-seq, we required highly enriched populations of mitotic cells. We used HeLa cells because this is a well-characterized cell line that has been used for a number of comparable studies. We found that a double thymidine block to synchronize cells at the G1/S boundary, followed by incubation with the microtubule poison nocodazole from 8 to 13 h post-release and a mitotic “shake-off” step routinely enriched mitotic cells to over 90% (as determined by flow cytometry with DNA and MPM-2 staining; Figure S2A, B). We also confirmed that the level of H3T3ph was largely preserved in preparations of sonicated chromatin from mitotic cells compared to whole HeLa cell lysates (Figure S2C). We then carried out ChIP-seq for H3T3ph and H3K4me2/3 on asynchronously growing and mitotically-enriched cell populations. As expected, we recovered DNA from H3K4me2/3 ChIPs from both asynchronous and mitotic cells and from H3T3ph ChIPs from mitotic cells, but little DNA was recovered for H3T3ph from asynchronous cells and these samples were not sequenced.

In previous immunofluorescence studies, H3T3ph emerges in early mitosis on chromosome arms, and becomes enriched around centromeres as prometaphase progresses (Dai et al., 2005; Markaki et al., 2009). Consistent with this, alignment of H3T3ph ChIP-seq data from nocodazole-arrested mitotic cells with the human genome revealed strong enrichment in the centromeric regions of all chromosomes (Figure 2A; Figure S3A, B). As previously reported, H3K4me2/3 was found predominantly at promoters in both asynchronous and mitotic cells (Figure S3C). In asynchronous cells, H3K4me2/3 flanked the nucleosome-depleted regions (NDRs) at promoters of active genes but, in mitosis, we observed a clear loss of NDR signals as previously described (Figure 2B), presumably due to nucleosomes spreading into the NDRs following dissociation of transcription factors (Festuccia et al., 2019; Javasky et al., 2018; Kelly et al., 2010b; Liang et al., 2015; Oomen et al., 2019).

**Figure 2.**
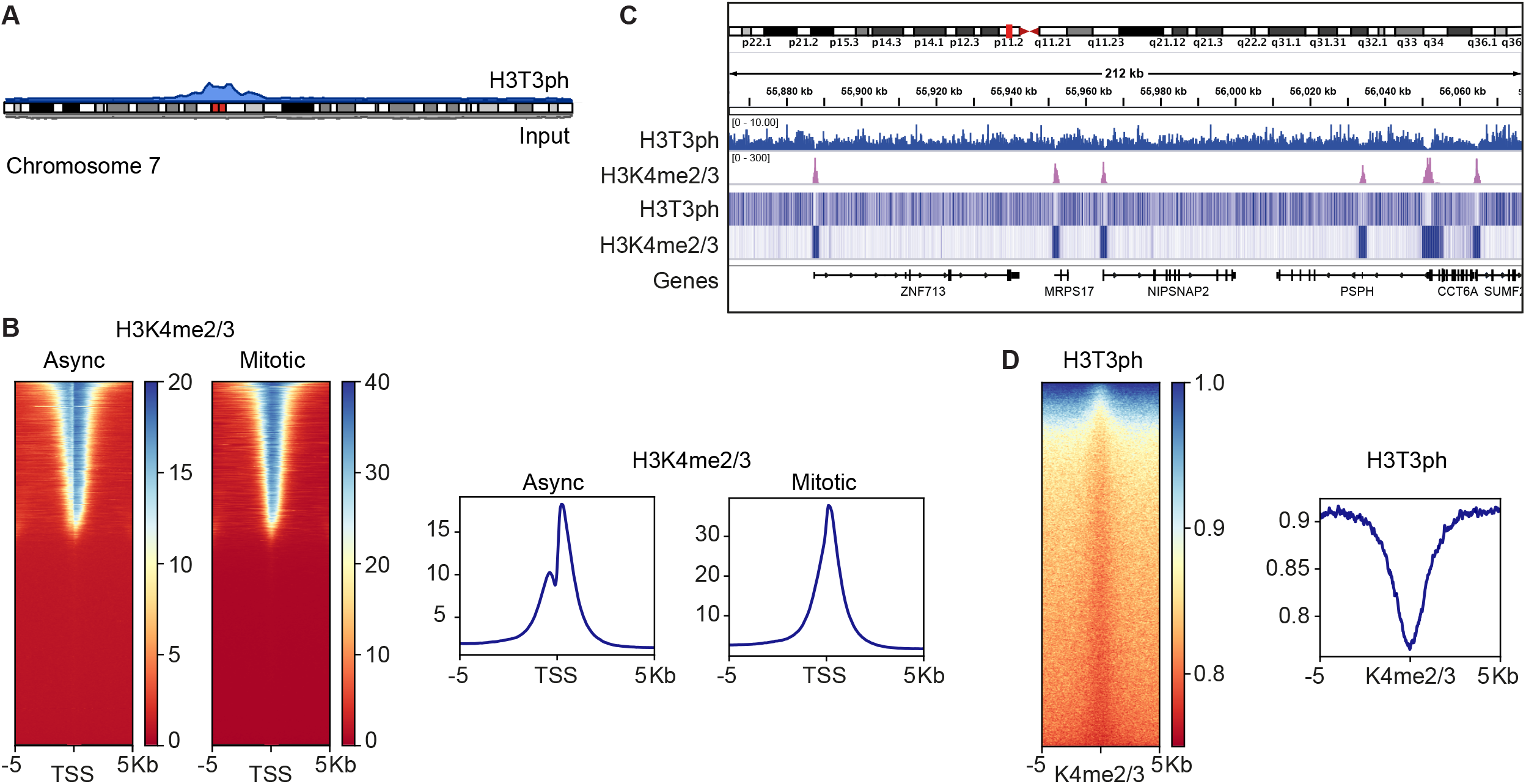
ChIP-seq for H3T3ph and H3K4me2/3 in HeLa cells. Asynchronous and mitotic HeLa cells were prepared as described in Figure S2. **A**. Ideogram showing enrichment of H3T3ph in mitosis at the centromere of human chromosome 7. H3T3ph (blue) and input (gray) read data from a single replicate were aligned to GRCh38.p12. Ideograms for all chromosomes are shown in Figure S3A. **B**. Genome-wide analysis of H3K4me2/3 enrichment at transcription start sites (TSS). Heatmaps (left) and metagene plots (right) showing H3K4me2/3 ChIP-seq of asynchronous and mitotic-enriched cells across 10 kb regions centered at TSSs. **C**. Integrative Genomics Viewer (IGV) tracks of mitotic H3T3ph and H3K4me2/3 ChIP-seq showing alignment on a 212 kb region of chromosome 7, represented by read coverage (input normalized) (middle panel) or heatmaps (bottom panel). **D**. Heatmap (left) and metagene plot (right) of H3T3ph ChIP-seq at 10 kb regions centered on mitotic H3K4me2/3 ChIP-seq peaks. Representative replicate data are shown in Figure S3B,C.

Although H3T3ph was enriched around centromeres, regions containing high levels of H3T3ph extended into euchromatic regions containing genes with H3K4me2/3 at their promoters. Strikingly, H3T3ph appeared to be selectively depleted at such promoters (Figure 2C). Indeed, when we examined the enrichment of H3T3ph around H3K4me2/3 peaks genome-wide in mitosis, the phosphorylation mark clearly declined in these regions (Figure 2D). These results suggest that the presence of H3K4me2/3 prevents the deposition of H3T3ph on adjacent sites in cells in mitosis.

### Phosphorylation of H3T3 is inhibited on nucleosomes carrying H3K4me3

A simple explanation for the absence of H3T3ph on mitotic chromatin in the vicinity of H3K4me2/3 would be if these methylation marks hinder the activity of H3T3 kinases. Indeed, previous work from our laboratory, and others, has shown that the activity of the mitotic H3T3 kinase Haspin on H3 peptides is progressively inhibited by mono, di and trimethylation of H3K4 (Eswaran et al., 2009; Han et al., 2011; Karimi-Ashtiyani and Houben, 2013). We confirmed that this was also true for H3 in nucleosomes, where H3K4me3 caused a clear reduction in the ability of Haspin to phosphorylate H3T3ph (Figure 3A).

**Figure 3.**
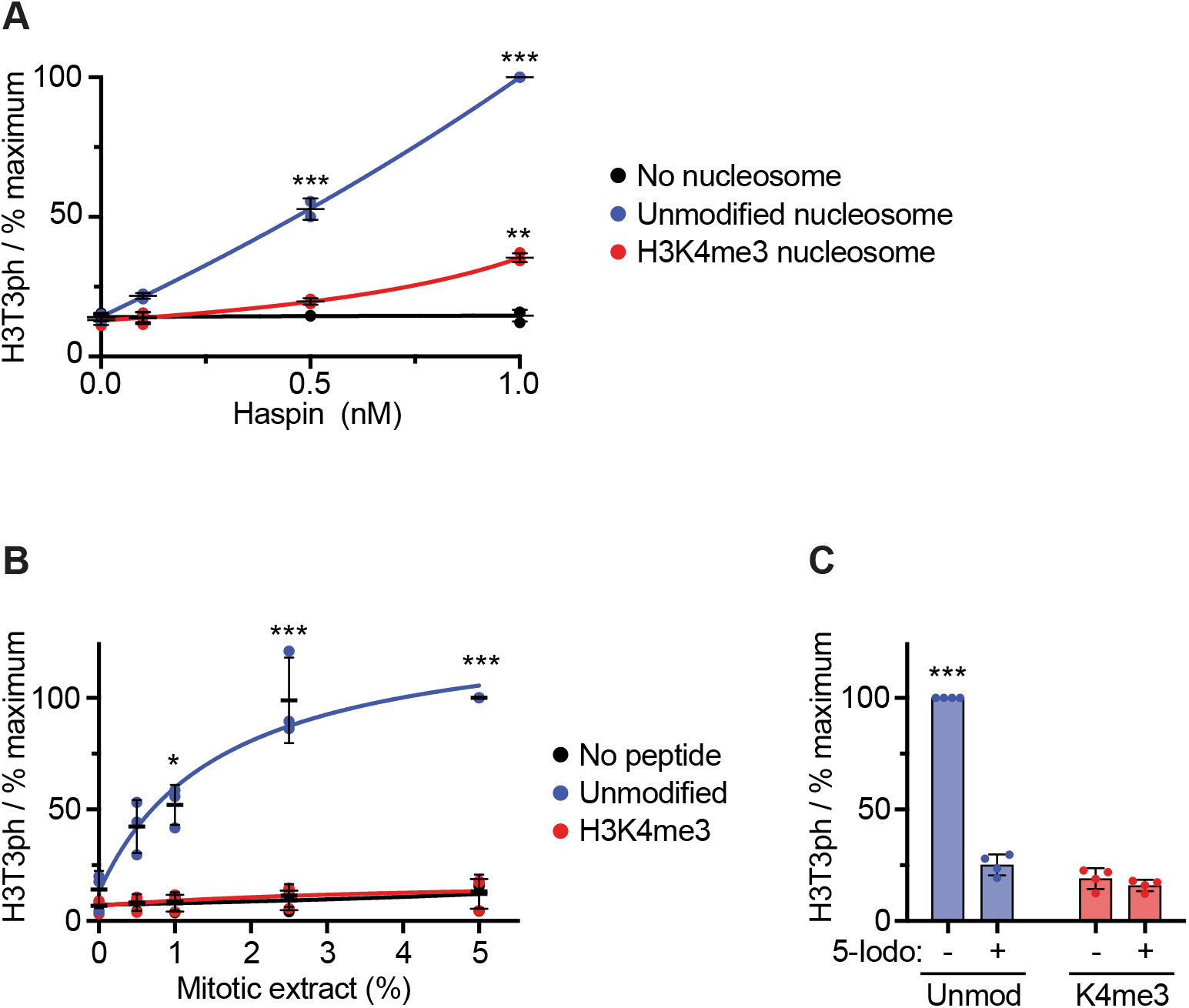
Haspin kinase activity towards H3T3 is inhibited by H3K4me3. **A**. *In vitro* kinase assay of recombinant human Haspin using recombinant nucleosomes with the indicated modifications as substrates (n = 3). **B**. Phosphorylation of modified H3 peptides driven by mitotic HeLa cell extracts (n = 3). **C**. Phosphorylation of modified H3 peptides driven by mitotic HeLa cell extracts in the presence and absence of the Haspin inhibitor 5-iodotubercidin (n = 4). Bars represent mean ± SD. Statistical analysis was carried out using non-normalised data, *** p < 0.0001, ** p < 0.001, * p < 0.01, when compared to phosphorylation in the absence of peptide.

Although there is strong evidence that Haspin is the major H3T3 kinase in mitotic cells (Cartwright et al., 2022; Dai et al., 2005; De Antoni et al., 2012; Markaki et al., 2009; Wang et al., 2012; Zhou et al., 2017), other potential kinases have been proposed (Kang et al., 2007; Karakkat et al., 2019). However, we found that phosphorylation of H3T3 driven by all endogenous kinases active in lysates of mitotic HeLa cells was severely curtailed by H3K4me3 (Figure 3B). Furthermore, the detected H3T3 kinase activity was strongly inhibited by a Haspin inhibitor, and so was ascribable to Haspin (Figure 3C). We conclude, therefore, that mitotic kinases are unable to efficiently phosphorylate H3T3 when the adjacent H3K4 is trimethylated.

### Haspin is not required to displace TFIID from mitotic chromatin

Previous work has shown that the PHD finger of TAF3 binds to H3K4me3, and to a lesser extent to H3K4me2 (Kungulovski et al., 2016; Shanle et al., 2017; van Nuland et al., 2013; Vermeulen et al., 2007). The binding sites of TAF3, and the TAF3 PHD finger alone, correlate well with the positions of H3K4me3 across the genome in asynchronous cells, suggesting that H3K4 methylation plays an important role in bringing TAF3 to chromosomes (Lauberth et al., 2013; Liu et al., 2011; Vermeulen et al., 2007). In experiments using synthetic H3 peptides *in vitro*, detectable binding of the TAF3 PHD, and of the TFIID complex containing TAF3, to H3K4me3 is eliminated by adjacent H3T3ph (see red box in Figure S4A) (Gatchalian et al., 2016; Kungulovski et al., 2016; Shanle et al., 2017; Varier et al., 2010). However, if H3T3ph generated by Haspin cannot be deposited adjacent to H3K4me2/3 in cells, then methyl-phos switching is unlikely and displacement of TFIID from chromosomes in mitosis should not be influenced by loss of H3T3ph. To test this, we examined the localization of TFIID components in cells lacking Haspin activity. All major TFIID components co-purify with GFP-TAF5 (van Nuland et al., 2013); multiple TAF proteins, including TAF5, are pulled down from cell lysates by H3K4me3-containing peptides (Vermeulen et al., 2007); and TAF5 has previously been used to monitor TFIID binding to H3K4me3 (Varier et al., 2010), making it a suitable choice for these experiments.

In immunofluorescence experiments using wild type HeLa cells, GFP-TAF5 was nuclear in interphase, and rapidly displaced from chromosomes in prophase, as previously reported (Varier et al., 2010) (Figure 4A). Indistinguishable results were obtained in HeLa cells that lacked detectable H3T3ph due to CRISPR-Cas9-mediated disruption of the Haspin gene (Figure 4A). To quantify these results, and to ensure they were not influenced by formaldehyde fixation artefacts (Teves et al., 2016), we analyzed the location of GFP-TAF5 in live cells throughout mitosis. Removal of GFP-TAF5 from chromosomes began within 5 min of nuclear envelope breakdown, and recovery started approximately 5 min after anaphase onset (Figure 4B, C; Figure S4B). This finding was consistent with multiple reports of TAF protein dissociation from mitotic chromosomes using a variety of methods, both with and without the use of formaldehyde (Djeghloul et al., 2020; Ginno et al., 2018; Samejima et al., 2022; Segil et al., 1996; Varier et al., 2010).

**Figure 4.**
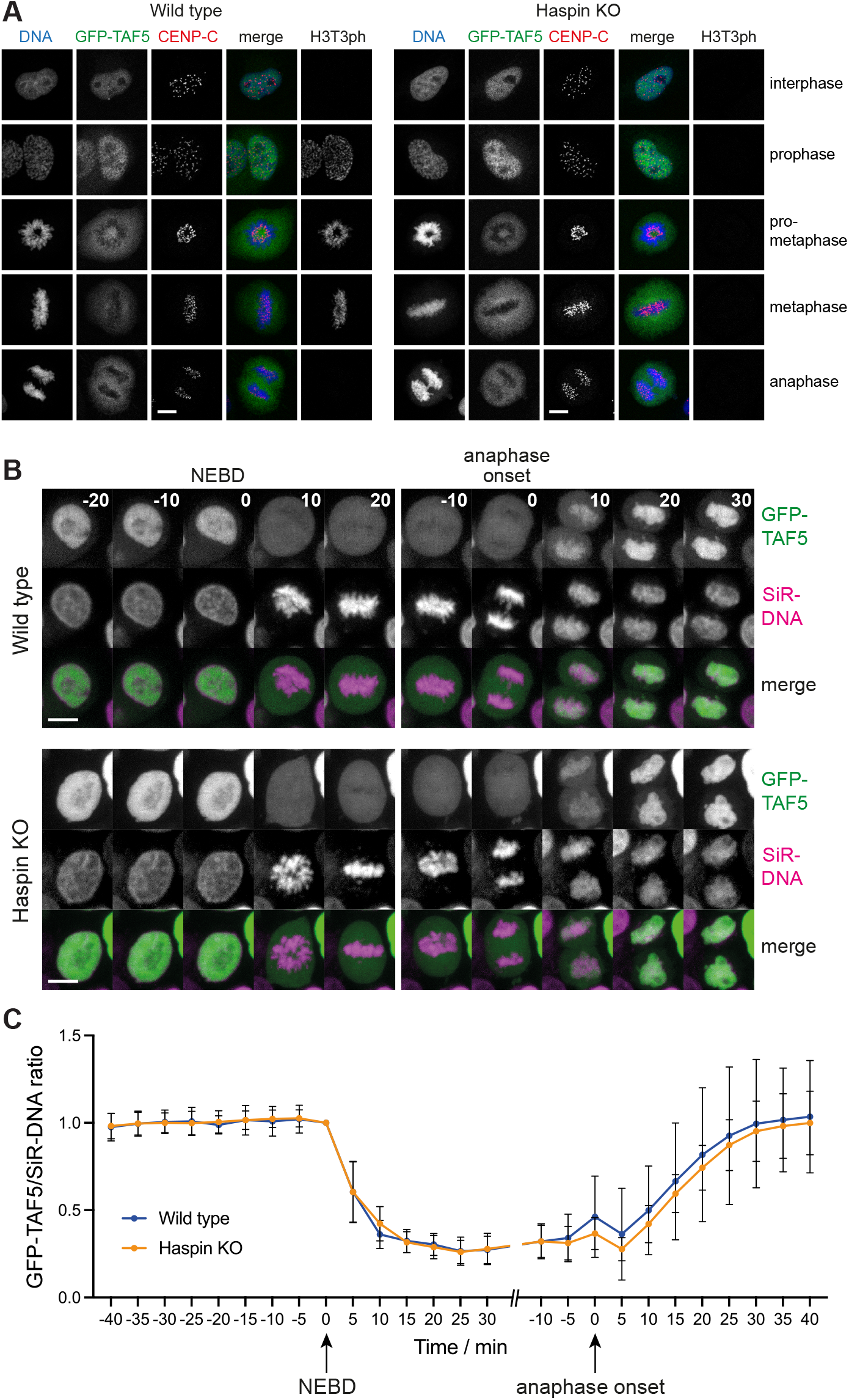
Haspin knock out does not influence the displacement of GFP-TAF5 from chromosomes in mitosis. **A**. Immunofluorescence microscopy (with formaldehyde fixation) for DNA (blue), GFP-TAF5 (green), CENP-C (centromeres, red), and H3T3ph (gray) in wild type and Haspin knockout HeLa cells. **B**. Live imaging of GFP-TAF5-expressing wild type and Haspin knockout HeLa cells. DNA was stained with SiR-DNA, images were taken every 5 min, and times are stated in minutes before and after nuclear envelope breakdown (NEBD) and anaphase onset as appropriate. For display purposes only, the intensities of GFP and SiR-DNA images were adjusted separately for each cell to allow visual comparison. Scale bars = 10 μm. **C**. Quantification of GFP-TAF5/SiR-DNA ratio during mitosis in live wild type (n = 9) and Haspin knockout (n = 10) HeLa cells imaged as in **B**, from 3 separate experiments. Bars represent mean ± SD. Individual cell traces are shown in Figure S4B. See also Figure S4C, and Videos 1 and 2.

Importantly, there was no significant difference in the results in wild type or Haspin knockout cells (two-way mixed effects model Anova, p = 0.30). When Haspin kinase activity was inhibited with 5-iodotubercidin (De Antoni et al., 2012; Wang et al., 2012), we also failed to see any change in GFP-TAF5 displacement by immunofluorescence microscopy in U2OS cells (Figure S4C), and similar results were obtained for GFP-TAF3 in HeLa cells (Figure S5). Finally, immunofluorescence for endogenous TAF3 gave indistinguishable results in wild type and Haspin knockout HeLa cells (Figure 5). Therefore, we could not reveal any effect of Haspin activity or H3T3ph on displacement or re-association of TFIID with chromosomes in mitosis.

**Figure 5.**
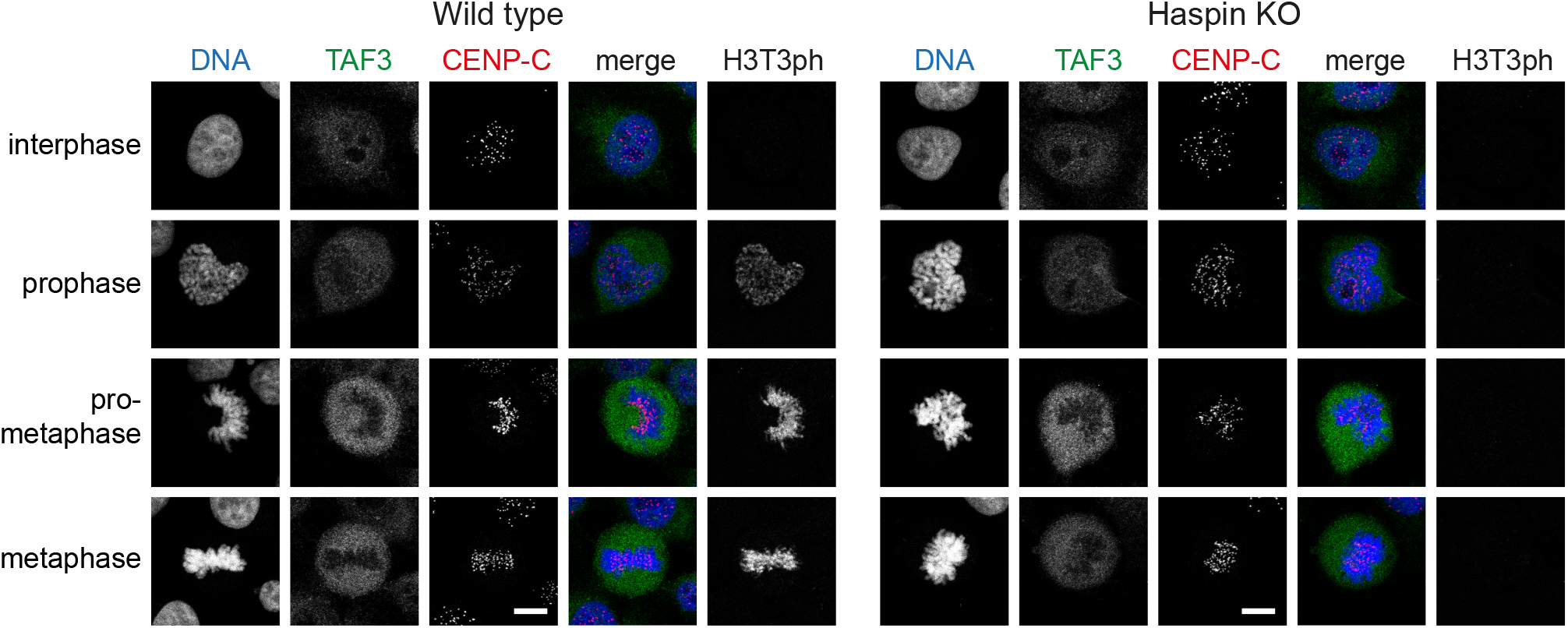
Haspin knockout does not influence the displacement of endogenous TAF3 from chromosomes in mitosis. Immunofluorescence microscopy (with formaldehyde fixation) for DNA (blue), TAF3 (green), CENP-C (centromeres, red), and H3T3ph (gray) in wild type and Haspin knockout HeLa cells. Scale bars = 10 μm. See also Figure S5.

### The TAF3 PHD finger binds similarly to interphase and mitotic chromatin

The results described above reveal the existence of an H3T3ph-independent mechanism to release TFIID from chromosomes in mitosis, which is likely to involve the phosphorylation of TFIID components by mitotic kinases (Segil et al., 1996). However, this does not rule out that a methyl-phos switch involving H3T3ph acts redundantly to modulate TAF3 binding to H3K4me2/3. To address this question, we adopted an approach in which we could determine the chromosome binding sites of the TAF3 PHD finger in the absence of other TFIID components and their post-translational modifications: chromatin interacting domain precipitation and sequencing (CIDOP-seq) (Kungulovski et al., 2016). In this variation of the ChIP-seq method, rather than using an antibody as the probe, we determined the genome-wide binding sites of a recombinant GST-TAF3 PHD finger fusion protein and, as a control, a M880A mutant that is defective in H3K4me2/3 binding (Kungulovski et al., 2016; Vermeulen et al., 2007). Notably, this approach also tests the influence of H3T3ph on TAF3 function in a way that does not rely on knowledge of histone antibody specificity.

We first confirmed that the GST-TAF3 PHD, but not the M880A mutant, was able to bind to H3K4me3-containing peptides *in vitro*, and that this binding was strongly blocked by adjacent H3T3ph (Figure S6A). As expected, GST-TAF3 PHD (but not the M880A mutant) pulled down H3K4me3-containing histone H3 from sheared chromatin of asynchronous HeLa cells as determined by immunoblotting (Figure S6B).

When using chromatin from mitotic cells, GST-TAF3 PHD (but not the M880A mutant) pulled down H3K4me3-containing H3 in similar amounts, and this did not contain H3T3ph (Figure S6B). Therefore, GST-TAF3 PHD can bind to mitotic chromatin, presumably at H3K4me3 sites. The presence of H3T3ph on mitotic chromatin did not detectably reduce GST-TAF3 PHD binding, likely because H3T3ph is not found adjacent to H3K4me2/3 in cells. However, from these results, we could not exclude that H3T3ph was able to prevent GST-TAF3 PHD binding to a subset of H3K4me3-containing regions, such as those surrounding centromeres.

To analyze this in further detail, we carried out CIDOP-seq to identify the genome-wide binding sites of the TAF3 PHD finger. As previously reported, TAF3 PHD bound to the chromatin of asynchronous cells in a pattern similar to that of H3K4me2/3, flanking the NDRs at promoters of active genes (Kungulovski et al., 2016) (Figure 6A, async; Figure S7A, B). As expected, M880A mutant TAF3 PHD showed only weak enrichment at H3K4me2/3 sites (Figure S7C). When using mitotic chromatin, GST-TAF3 PHD retained the ability to bind promoters and spread into NDRs, consistent with the incursion of nucleosomes (Figure 6A, mitotic), as seen for H3K4me2/3 ChIP-seq (see Figure 3). Importantly, TAF3 PHD was equally enriched at transcription start sites of genes in centromere proximal regions with high overall levels of H3T3ph (Figure 6B) as it was genome wide (Figure 6A), or along chromosome arms where H3T3ph was low (Figure S7D). These results confirmed that H3T3ph does not prevent the TAF3 PHD interaction with H3K4me3 on mitotic chromosomes. Therefore, we conclude that, although H3T3ph does displace TAF3 from H3K4me3 *in vitro*, the absence of H3T3ph immediately adjacent to H3K4me2/3 in cells precludes the operation of a methyl-phos TAF3 switch *in vivo*.

**Figure 6.**
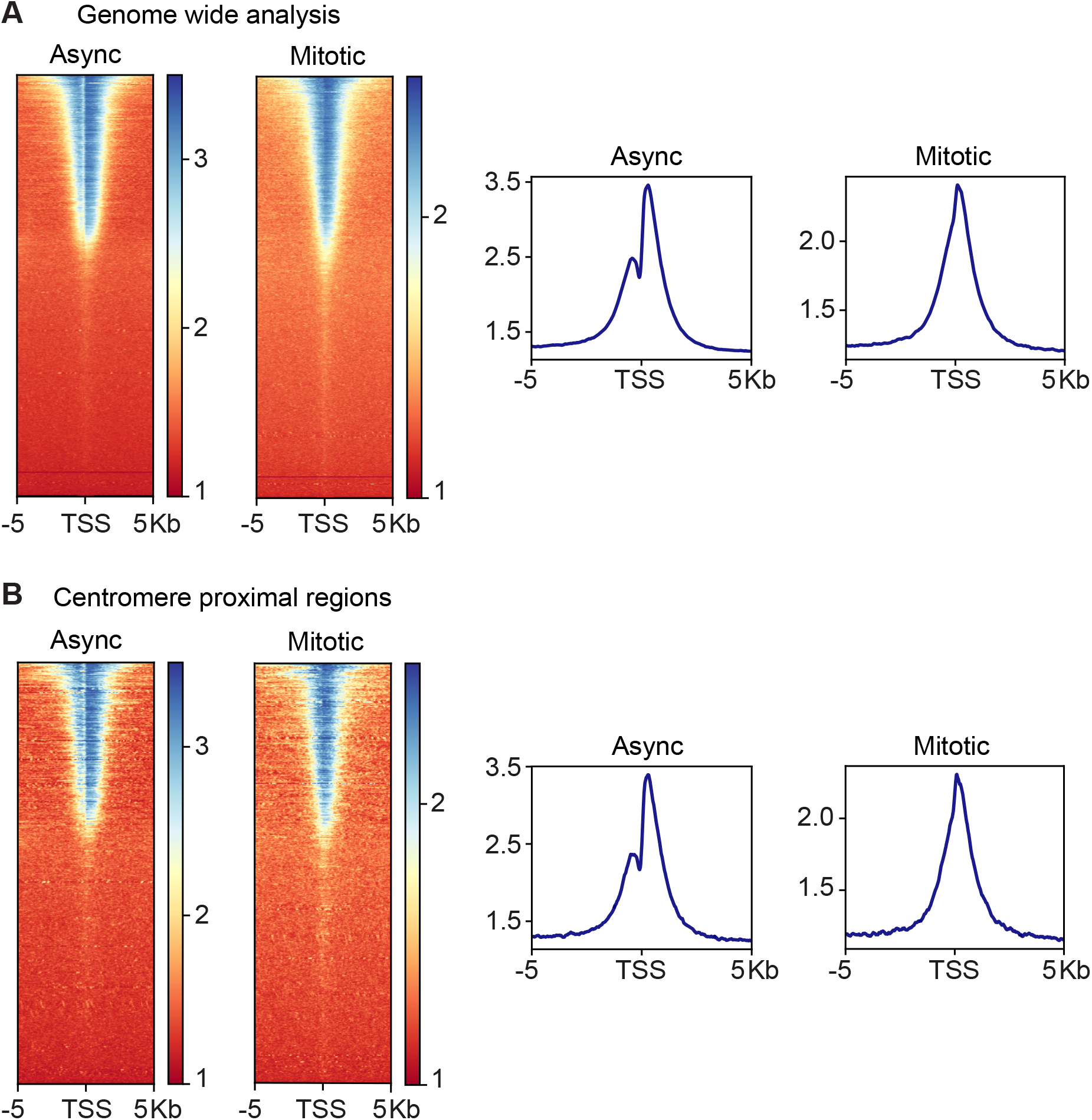
CIDOP-seq shows TAF3 PHD binding to interphase and mitotic chromatin. **A**. Genome wide analysis of TAF3 PHD enrichment at TSSs. **B**. TAF3 PHD enrichment at centromere proximal TSSs. Heatmaps (left) and metagene plots (right) show TAF3-PHD binding across 10 kb regions centered at TSSs for both asynchronous and mitotic-enriched HeLa cells. See also Figures S6 and S7.

### Absence of a methyl-phos switch for additional H3K4 reader proteins

Finally, we wanted to test if other reported H3K4 reading proteins are influenced by methyl-phos switching with H3T3ph. ING2 and Dido3 are H3K4me3-reading proteins that no longer bind H3K4me3 *in vitro* if H3T3ph is adjacent (Garske et al., 2010; Gatchalian et al., 2013; Jain et al., 2020; Tencer et al., 2017; Varier et al., 2010). As previously reported (Gatchalian et al., 2013; Gatchalian et al., 2016; Trachana et al., 2007), ING proteins and Dido3 were displaced from chromosomes in mitosis and we found that this was indistinguishable in cells lacking Haspin and H3T3ph (Figure 7A; Figure S8A).

**Figure 7.**
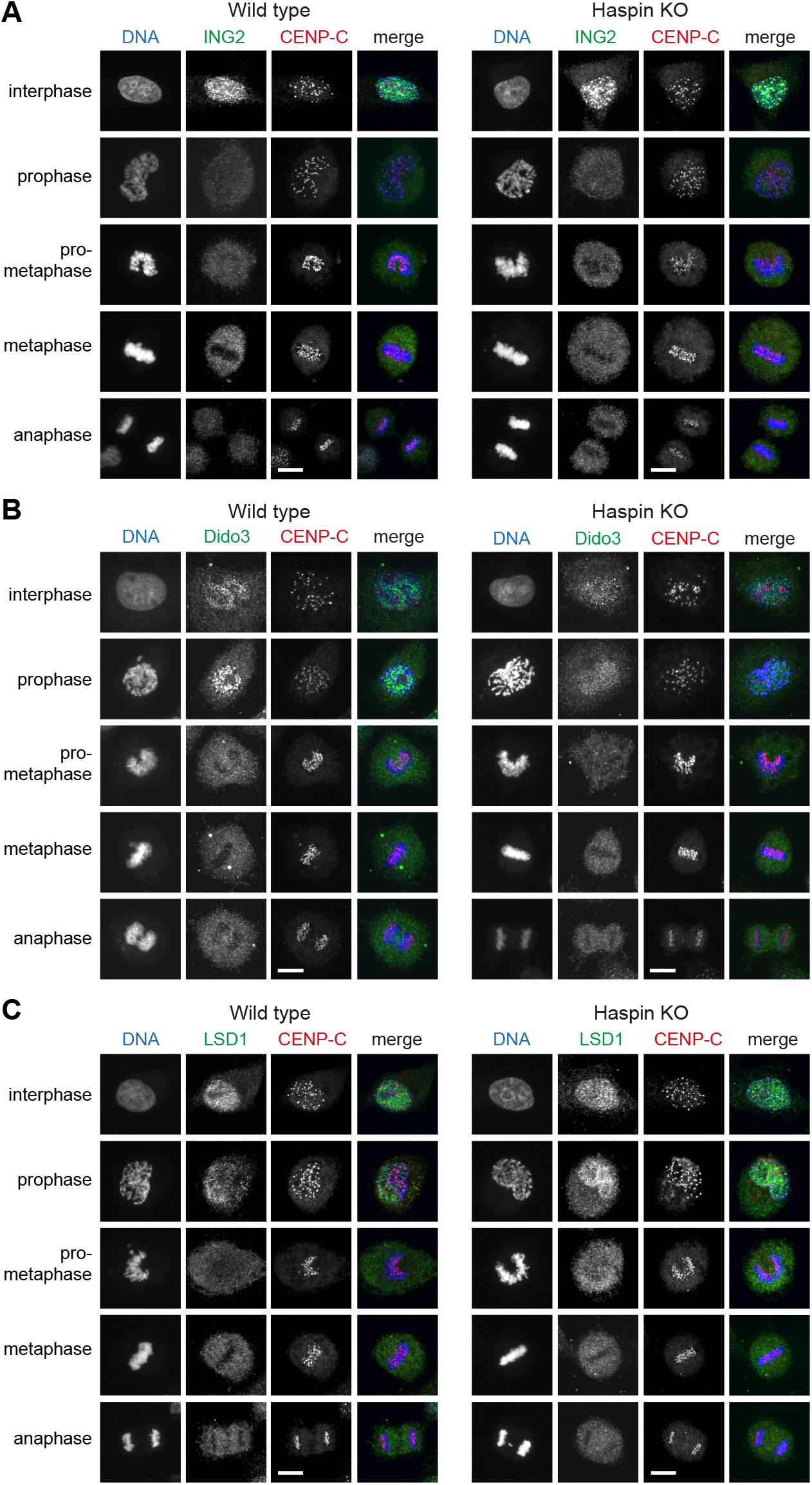
Haspin knockout does not influence the displacement of endogenous ING2, Dido3, or LSD1 from chromosomes in mitosis. **A**. Immunofluorescence microscopy (with methanol fixation) for DNA (blue), ING2 (green), CENP-C (centromeres, red), and H3T3ph (gray) in wild type and Haspin knockout HeLa cells. **B**. As for A, but staining for Dido3 (green). **C**. As for A, but staining for LSD1 (green). Scale bars = 10 μm. See also Figure S8.

Interestingly, the displacement of the H3K4me0-reading LSD1 complex from mitotic chromosomes was also not influenced by loss of H3T3ph (Figure 7B; Figure S8B), even though detectable binding of the LSD1 complex component BHC80 to H3 peptides is eliminated by H3T3ph *in vitro* (Garske et al., 2010; Varier et al., 2010). We were, therefore, unable to observe methyl-phos switching at H3T3-H3K4 in cells.

## DISCUSSION

Methyl-phos switching is an appealing concept that has been invoked to explain the dissociation of numerous proteins from chromosomes during cell division. In the case of factors that read the modification state of H3K4, more than thirty proteins have been found to be displaced by H3T3ph *in vitro*, and this has been proposed to be important *in vivo* (see Introduction). Our results, however, suggest that H3T3ph is rarely deposited next to H3K4me2/3 in cells. Therefore, the displacement of H3K4me2/3 reader proteins from mitotic chromatin is unlikely to be due to methyl-phos switching (see Figure S4A).

Previous studies from our group and others have shown that the kinase activity of human and *Arabidopsis* Haspin towards H3T3 within histone peptides is progressively inhibited by mono-, di- and tri-methylation of H3K4 (Eswaran et al., 2009; Han et al., 2011; Karimi-Ashtiyani and Houben, 2013). Indeed, the crystal structure of Haspin in complex with an H3 substrate peptide suggests that methylation of H3K4 would interfere with phosphorylation of H3T3 (Maiolica et al., 2014). We extend this here, showing that H3K4me3 substantially blunts the ability of human Haspin (and, indeed, any kinase active in mitotic cell lysates) to deposit H3T3ph on nucleosomes *in vitro*. Therefore, we conclude that the diminished H3T3ph in chromatin regions containing H3K4me2/3 is due to blocking of Haspin activity at these sites (Figure S4A). Interestingly, a dip in H3T3ph at active promoters containing H3K4me3 has also been observed in *Arabidopsis* and possibly in *Chlamydomonas*, and H3T3ph is reduced in promoter regions containing active RNA polymerase II (Pol II S5ph) in *Drosophila* S2 cells (Casas-Mollano et al., 2008; Fresan et al., 2020; Wang et al., 2015). Although the kinase depositing H3T3ph and/or the cell cycle stage involved was less clear in these studies, the antagonistic effect of H3K4me2/3 on H3T3ph appears to be conserved.

A previous study suggested the existence of a combinatorial mark on histone H3 in mitosis, comprising H3T3ph, H3K4me3 and H3R8me2 (Markaki et al., 2009). However, it is difficult to obtain definitive evidence for a particular combination of marks using polyclonal mixtures of antibodies or MALDI-TOF alone (Bonaldi et al., 2004; Soupsana et al., 2021). Of course, we cannot rule out that adjacent H3T3ph and H3K4me2/3 marks are present at a low level, but tandem mass spectrometry studies of histone modifications to date have found H3T3ph alone or in combination with H3K4me1, but not with H3K4me2/3 (Bonenfant et al., 2007; Garcia et al., 2005; Jung et al., 2013; Maile et al., 2015), consistent with our ChIP-seq data.

The results of prior work appeared to support the idea that Haspin and H3T3ph regulate the association of TAF3 with H3K4me3 in cells (Varier et al., 2010). A significant finding in this previous study was that overexpression of Haspin could repress TAF3-mediated gene activation. However, Haspin is many-fold overexpressed in these experiments, leading to increased H3T3 phosphorylation in both interphase and mitosis (Dai et al., 2005; Varier et al., 2010). Because high concentrations of Haspin can overcome the suppressive effect of H3K4me3 *in vitro*, and we know that H3T3ph does displace TAF3 from H3K4me3 peptides *in vitro* (Gatchalian et al., 2016; Kungulovski et al., 2016; Shanle et al., 2017; Varier et al., 2010), it seems likely that elevating Haspin activity was sufficient to artificially flip the methyl-phos switch in this previous work. It is less clear why Haspin RNAi led to GFP-TAF5 retention on mitotic chromosomes in the previous study, in contrast to our current results using both Haspin inhibitor and knockout cells.

Some previous cell-based studies provided only indirect evidence for H3T3ph/H3K4me3 switching because they relied on the inverse correlation of H3T3ph and H3K4me3 reader protein levels on chromosomes (Ali et al., 2013; Gatchalian et al., 2013). Our results are not at odds with the idea that a number of H3K4me-reading proteins are lost from chromosomes in mitosis, but they argue that H3T3ph is not responsible. Treatment with the Haspin inhibitor CHR-6494 decreased H3T3ph in another report, and this was suggested to cause retention of the H3K4me3-reading proteins Dido3 and ING1 on chromatin during mitosis (Gatchalian et al., 2016). However, there was little detectable difference in reader dissociation from chromosomes as cells entered mitosis in the absence of H3T3ph in this study. It was proposed instead that there was more rapid re-recruitment of Dido3 and ING1 to chromosomes in telophase, but the difficulty in distinguishing different stages of late mitosis using fixed cell images, and the use of a Haspin inhibitor whose selectivity is poorly characterized (Huertas et al., 2012), means that care must be taken in interpreting these qualitative results. By contrast, we find no change in the loss and re-recruitment of TAF5, Dido3, and ING2 to chromosomes during mitosis using live and fixed cell imaging, either in cells genetically lacking Haspin, or in cells where Haspin was inhibited by the well-characterized inhibitor 5-iodotubercidin, in which H3T3ph is undetectable (De Antoni et al., 2012; Wang et al., 2012; Zhou et al., 2017). Instead, direct phosphorylation of transcription factors is likely to be required for mitotic displacement (Figure S4A; Gottesfeld and Forbes, 1997; Segil et al., 1996; Segil et al., 1991).

The technical difficulties inherent in determining if crosstalk between histone marks occurs in cells are well-known (Fischle, 2008; Rothbart et al., 2015; Shah et al., 2018). Our study provides an example of an effective set of approaches to this general problem. Importantly, we used CIDOP-seq to confirm that the PHD finger of TAF3 is able to bind to sites containing H3K4me2/3 in both mitotic and interphase chromatin, even in regions near centromeres that have the highest overall levels of H3T3ph in mitosis. This approach may be useful for analyzing the function of other combinatorial histone marks without the uncertainties introduced by the use of antibodies that inevitably have incompletely defined specificity for naturally modified histones.

What might be the significance of excluding H3T3ph from the vicinity of H3K4me3? One possibility is that it allows H3K4me2/3 function and promoter identity to be maintained during mitosis, perhaps including a basal level of mitotic transcription. Absence of H3T3ph could prevent the recruitment and functional impact of H3T3ph-dependent reader proteins at the promoters of active genes in mitosis. For example, Aurora B, which is recruited by H3T3ph as part of the CPC, can alter transcription factor activity (Frangini et al., 2013; Kassardjian et al., 2012; Shin et al., 2016; Wu et al., 2011), which might need to be avoided during mitosis. Exclusion of H3T3ph could also permit H3K4me2/3 functions in mitosis, unhindered by competition from adjacent H3T3ph-binding proteins or methyl-phos switching activity. Indeed, although most H3K4me2/3 readers are largely displaced in mitosis, a subset of promoters may retain H3K4me2/3 reading proteins (Christova and Oelgeschlager, 2002; Segil et al., 1996). In addition, condensins (which facilitate chromosome compaction in mitosis) appear to be loaded onto chromatin at active promoters (Dowen et al., 2013; Kim et al., 2013; Sutani et al., 2015; Toselli-Mollereau et al., 2016) and, for Condensin II, this may be facilitated by recognition of H3K4me3 (Yuen et al., 2017).

Our results leave open the possibility that some readers of unmethylated or monomethylated H3K4 might still be regulated by H3T3ph. Other work has suggested that mutation of the H3K4me0 reader DNMT3A, to render it insensitive to H3T3ph, increases its association with H3T3ph-containing regions of mitotic chromosomes (Noh et al., 2015). These results provide initial evidence to support the importance of H3T3ph in regulating the localization of H3K4me0 readers in cells. Another H3K4me0 reader, PHF21A/BHC80 is strongly displaced from H3 peptides by H3T3ph *in vitro* (Garske et al., 2010; Varier et al., 2010). However, we were unable to detect any effect of Haspin knockout on the displacement of the PHF21A/BHC80-containing LSD1 complex from mitotic chromosomes. Our findings highlight that, regardless of the results of *in vitro* binding assays, the effect of combinations of histone marks on the binding of histone reading proteins must be examined in cells on a case-by-case basis before conclusions can be drawn about functional significance. A H3T3-H3K4me2/3 methyl-phos switch can be created *in vitro*, but our results suggest this mechanism does not function *in vivo* and is unlikely to be involved in widespread transcriptional repression in cells.

## AUTHOR CONTRIBUTIONS

**RJH:** Conceptualization, Investigation, Formal analysis, Visualization, Writing - Original Draft. **MH:** Software, Formal analysis. **MDL:** Investigation. **TNC:** Investigation. **BW:** Investigation. **JLM:** Formal analysis. **JMC:** Investigation, Methodology. **LG:** Methodology. **LP:** Investigation, Methodology. **DR:** Conceptualization, Writing - Review & Editing, Supervision, Funding acquisition. **JMGH:** Conceptualization, Formal analysis, Visualization, Writing - Original Draft, Supervision, Project administration, Funding acquisition.

## ACKNOWLEDGMENTS

We would like to sincerely thank the late Dr Robert Stones, of the Newcastle University Bioinformatics Support Unit, for his contributions to the early stages of this work. We also thank Dr Albert Jeltsch (Institute of Biochemistry and Technical Biochemistry, University of Stuttgart) for providing constructs encoding the TAF3 PHD finger, Dr Marc Timmers (German Cancer Research Center, University of Freiburg) for constructs and cell lines expressing GFP-TAF3 and TAF5, and Dr Fangwei Wang (Life Sciences Institute, Zhejiang University) for Haspin knockout cells. The authors gratefully acknowledge the Newcastle University BioImaging Unit, Genomics Core Facility, Bioinformatics Support Unit, and Flow Cytometry Core Facility (FCCF) for their support and assistance in this work. This study was funded by a Wellcome Trust Investigator Award (106951/Z/15/Z) and a Royal Society Wolfson Research Merit Award (WM130089) to JMGH, an EPSRC DTP (Biological Informatics) PhD Studentship to MH, a Barbour Foundation PhD Studentship to JLM, and by a JGW Patterson Foundation grant to LG. For the purpose of open access, the author has applied a CC BY public copyright license to any Author Accepted Manuscript version arising from this submission.

## METHODS

### Antibodies

| Antibody | Source | IF dilution | IB dilution | ELISA dilution | ChIP (μg) | Flow cytometry | RRID |
| --- | --- | --- | --- | --- | --- | --- | --- |
| Rabbit pAb Histone H3T3ph (B8634) | (Dai et al., 2005) | 1/1000 | 1/500 | 1/4000 | 8 |  |  |
| Rabbit mAb Histone H3K4me3 (C42D8) | Cell Signaling Technology #9751 |  | 1/500 | 1/100,000 | 3 |  | RRID:AB_2616028 |
| Rabbit pAb Histone H3 | Abcam ab1791 |  | 1/100,000 |  |  |  | RRID:AB_302613 |
| Chicken pAb GFP | Abcam ab13970 | 1/1000 |  |  |  |  | RRID:AB_300798 |
| Guinea pig pAb CENP-C | MBL PD030 | 1/2000 |  |  |  |  | RRID:AB_10693556 |
| Mouse mAb TAF3 (39TA-2F5) | Invitrogen MA3-074 | 1/250 |  |  |  |  | RRID:AB_2633320 |
| Rabbit pAb Dido/DATF1 | Invitrogen PA5-101330 | 1/200 |  |  |  |  | RRID:AB_2850767 |
| Rabbit pAb ING2 | Sigma HPA019486 | 1/500 |  |  |  |  | RRID:AB_1851780 |
| Rabbit pAb LSD1/KDM1A | Abcam ab17721 | 1/200 |  |  |  |  | RRID:AB_443964 |
| Mouse mAb GST (GST 3-4C) | Invitrogen 13-6700 |  | 1/5000 | 1/5000 |  |  | RRID:AB_2533028 |
| Mouse mAb Cy5-conjugated MPM2 antibody | Sigma 16-220 |  |  |  |  | 1/100 | RRID:AB_442398 |
| Goat anti-rabbit-IgG Alexa Fluor 488 | Invitrogen A-32731 | 1/1500 |  |  |  |  | RRID:AB_2633280 |
| Donkey anti-mouse-IgG Alexa Fluor 488 | Invitrogen A-21202 | 1/1500 |  |  |  |  | RRID:AB_141607 |
| Goat anti-chicken IgY Alexa Fluor 488 | Invitrogen A-21449 | 1/1500 |  |  |  |  | RRID:AB_2535866 |
| Donkey anti-rabbit IgG Alexa Fluor Plus 594 | Invitrogen A-32754 | 1/1500 |  |  |  |  | RRID:AB_2762827 |
| Goat anti-guinea pig IgG Alexa Fluor 647 | Invitrogen A-21450 | 1/1500 |  |  |  |  | RRID:AB_2735091 |
| Goat anti-rabbit IgG-HRP | Cell Signaling Technology #7074 |  | 1/1000 | 1/2000 |  |  | RRID:AB_2099233 |
| Horse anti-mouse IgG-HRP | Cell Signaling Technology #7076 |  | 1/1000 | 1/2000 |  |  | RRID:AB_330924 |

### Peptides

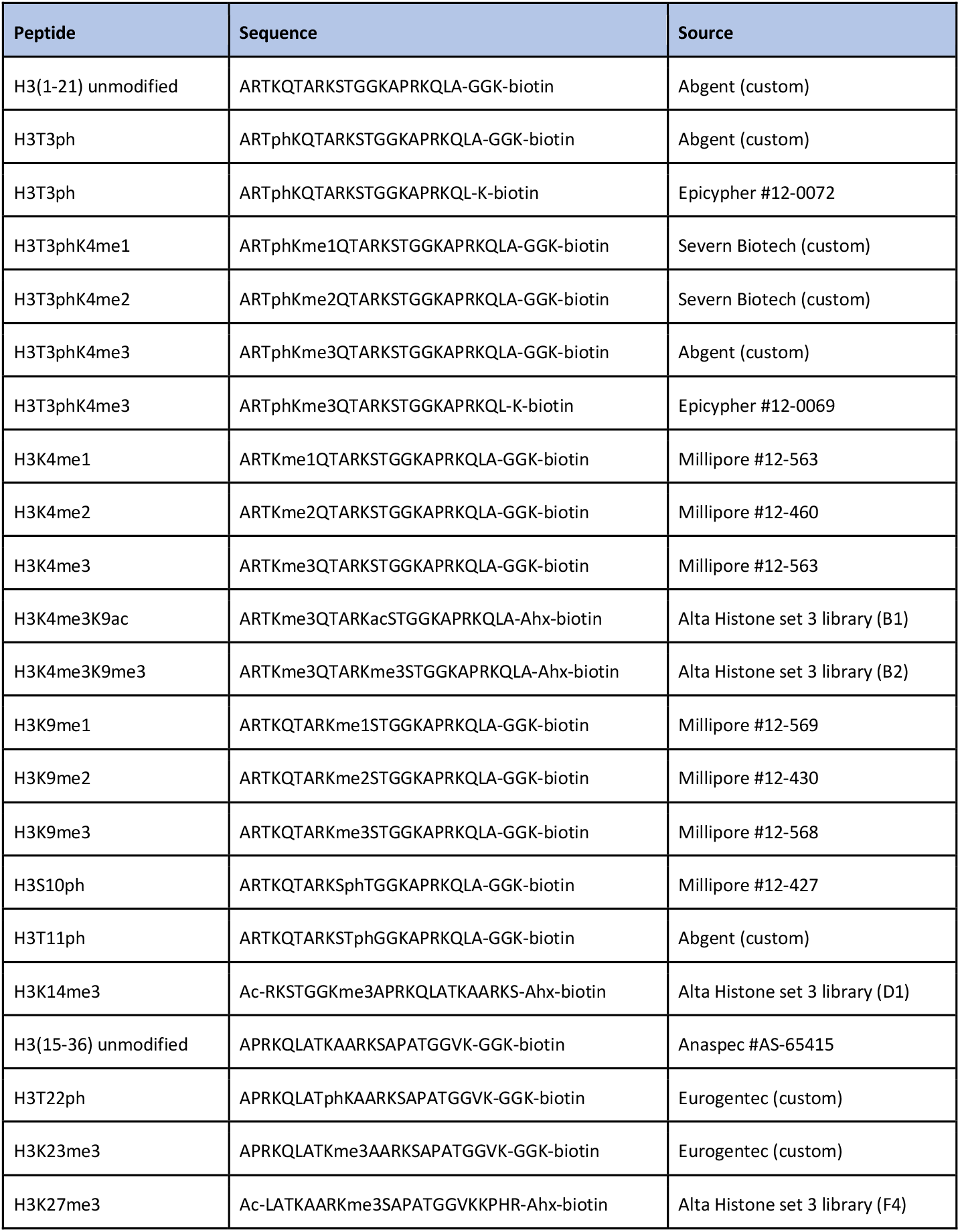

### Cell culture

HeLa S3 (ATCC CCL-2.2; RRID:CVCL_0058), parental and Haspin knockout D2 Hela cells (Zhou et al., 2017), human GFP-TAF5 U2OS cells (Varier et al., 2010), and doxycycline-inducible mouse GFP-TAF3 HeLa FRT (van Nuland et al., 2013) and human GFP-TAF5 HeLa FRT cells (Antonova et al., 2018) were maintained in high glucose DMEM with 10% (v/v) FBS, 100 U/ml penicillin and streptomycin and 2 mM L-glutamine at 37°C and 5% CO_2_ in a humidified incubator. Doxycycline inducible lines were induced for 24 h with 0.5 - 1 μg/ml doxycycline. Where stated, cells were treated with 1 μM 5-iodotubercidin for 2 h.

Parental and Haspin knockout cells stably expressing GFP-TAF5 were obtained by cotransfection with pcDNA5/FRT/TO/GFP-TAF5 (Antonova et al., 2018) and a pPGKpuro resistance plasmid (in a 9:1 ratio) using Xtreme Gene 9 (Roche). After 48 h, 2 μg/ml puromycin was added and, after an additional 10 days, cells expressing moderate levels of GFP-TAF5 were isolated by fluorescence-activated cell sorting.

### Cell synchronization

For ChIP-seq experiments, mitotic HeLa cells were removed from cultures at approximately 25% confluency by shake off before adding 2.5 mM thymidine to the remaining adherent cells for 18 h. Thymidine was removed for 8 h, and re-added for 18 h before an 8 h release into medium lacking thymidine. Nocodazole was added at 0.5 μM for 5 h and mitotic cells collected by gentle shake off.

To assess synchronization by flow cytometry, cells were fixed in ice cold 70% ethanol and permeabilized with 0.1% Triton-X100 in PBS for 5 min. After centrifugation, cells were resuspended in 0.1% BSA, PBS with 1/100 Cy5-conjugated MPM2 antibody for 1 h. Cells were then washed and resuspended in PI staining buffer (50 μg/ml propidium iodide and 0.2 mg/ml RNase A in PBS) and analysed on a FACSCanto II cell analyser (BD Biosciences) and FCS Express 7 (De Novo Software).

### Recombinant proteins and peptides

We used recombinant biotinylated human mononucleosomes (Epicypher 16-0006), H3K4me3 mononucleosomes (Epicypher 16-0316), and H3T3ph and H3T3phK4me3 mononucleosomes (Epicypher, custom). Recombinant human 6His-Haspin residues 471-798 was produced as previously described (Dai et al., 2005).

Plasmids encoding recombinant WT (RRID:Addgene_92100) and M880A mutant human GST-TAF3-PHD were kindly provided by Albert Jeltsch (Kungulovski et al., 2016). *E. coli* BL21 cells carrying each plasmid were grown in LB medium at 37°C in the presence of 100 μM ZnSO_4_ to an A_600_ of 0.6, then induced with 1 mM IPTG overnight at 20°C. Collected cells were resuspended in 20 mM HEPES, 500 mM KCl, 0.2 mM DTT, 1 mM EDTA, 10% glycerol, 1 mg/ml lysozyme, 1% Benzonase and cOmplete Protease Inhibitor Cocktail (Roche), pH 7.5, and disrupted by probe sonication. GST-TAF3 PHD proteins were purified using the Pierce GST Spin Purification Kit (ThermoFisher) and dialyzed against 20 mM HEPES, 200 mM KCl, 0.2 mM DTT, pH 7.5 with 10% glycerol for 2 h, then with 60% glycerol overnight.

### ELISAs and protein interaction assays

For characterization of antibody binding to histone peptides, Streptavidin Coated High Capacity 96 well plates (Pierce) were washed 3 times with TBST (TBS, 0.05% Tween20) before adding 100 μl 0.1 μM peptide in TBST. After incubation at room temperature for 1 - 2 h, wells were washed and 100 μl primary antibody in TBST, 0.1% BSA were added and incubated for 1 - 2 h. After washing, 100 μl 1/2000 HRP-conjugated anti-rabbit or anti-mouse IgG secondary antibodies (Cell Signaling Technology) were added for 1 h at room temperature before washing again. The signal was detected using TMB substrate (Pierce) according to the manufacturer’s instructions. Absorbance was measured at 450 nm (signal) and 570 nm (background) using a Polarstar Omega microplate reader (BMG Labtech). ELISAs using recombinant mononucleosomes were carried out in 384-well high capacity streptavidin-coated plates (Pierce).

Mononucleosomes were used at 0.1 μM in 40 μl TBST. Antibodies were used as above at 40 μl per well. For antibody binding assays in the presence of sheared chromatin, 0.8 μg H3T3ph antibody B8634 and 0.1 μM mononucleosomes were incubated with 20 μl sheared chromatin (obtained as for ChIP-seq), shaking at 300 rpm, for 2 h in a 384-well plate. Samples were then moved to a 384-well high capacity streptavidin-coated plate and antibody binding was detected by ELISA as just described.

Analysis of GST-TAF3-PHD binding to histone peptides (coated at 0.1 μM in 40 μl TBST) was carried out in 384-well high capacity streptavidin-coated plates (Pierce). GST-TAF3-PHD (WT or M880A mutant) were made to equivalent concentrations (∼0.1 mg/ml) in interaction buffer (20 mM HEPES, 300 mM KCl, 0.1 mM DTT, 10% glycerol, pH 7.5). After GST-TAF3-PHD binding to peptides for 1 h at room temperature, the plates were washed 3 times with interaction buffer, then once with TBST, before anti-GST antibody (Invitrogen) binding and detection as described above.

### *In vitro* kinase assays

Reactions contained recombinant 6His-Haspin, 0.1 μM recombinant biotinylated human mononucleosomes, and 0.2 mM ATP in K buffer (50 mM Tris, 10 mM MgCl_2_, 1 mM EGTA, 10 mM NaF, 20 mM β-glycerophosphate, 1 mM PMSF, pH 7.5 with PhosSTOP (Merck)), and were carried out in 384-well microplates at 37°C for 30 min. Samples were then moved to a 384-well high capacity streptavidin-coated plate, and phosphorylation detected using H3T3ph antibody (B8643) as described for ELISA.

For kinase assays using whole cell extract, HeLa cells were treated with 300 nM nocodazole for 15 h, collected by shake off, and lysed at 4°C in 50 mM Tris, 0.25 M NaCl, 0.1% Triton X100, 10 mM MgCl_2_, 2 mM EDTA, 1 mM DTT, pH 7.5, with protease inhibitor cocktail (Sigma P8340), PhosSTOP, 1 mM PMSF, 0.1 μM okadaic acid, 10 mM NaF and 10 mM β-glycerophosphate, at approximately 30 × 10^6^ cells/ml. Extracts were immediately flash frozen in liquid nitrogen. Kinase reactions contained 0.35 μM biotinylated peptide, 0.2 mM ATP and up to 5% mitotic extract in K buffer. Where indicated, 5-iodotubercidin was included at 10 μM. Phosphorylation was detected as described above.

### Immunofluorescence microscopy

Cells grown on glass coverslips or 8-well chamber slides (Eppendorf) were either fixed for 10 min with 4% (v/v) formaldehyde in PBS, washed twice with PBS, and permeabilised for 2 to 5 min with 0.1% Triton X100 in PBS, or were treated with ice cold methanol for 10 min. After washing twice with PBS, samples were blocked for 1 h with 1 to 5% BSA in PBS, 0.05% Tween 20 (PBST). Primary antibodies in blocking buffer were added for 1 - 2 h at 37°C then washed twice with PBST before incubating with fluorophore-conjugated secondary antibodies for approximately 1 h in blocking buffer. After washing twice with PBST, once with PBS, and once with H_2_O, mounting was carried out with ProLong Glass with NucBlue (ThermoFisher).

Confocal imaging was performed using a Leica SP8 confocal microscope with a 63x NA1.4 Plan Apo CS2 Oil objective using Leica LasX v3 software, or a Nikon A1 confocal microscope equipped with a 60x NA1.4 Apo Oil *λ*S DIC N2 objective using Nikon Elements 5.22 software, or a Visitech VT-iSIM super-resolution Nikon TiE-based microscope (Visitech, UK) with instant SIM scanhead coupled to two Hamamatsu Flash4 v2 cameras (Hamamatsu, Japan) via a dual port splitter, using a 60x NA1.4 Plan Apo VC Oil DIC N2 objective and Elements 5.21.03 software (Nikon, Japan). Images were processed in ImageJ 2.3.0 (Schindelin et al., 2015) and Adobe Photoshop 2022, and are displayed as maximum intensity projections of the central 5 slices.

### Live imaging

Live imaging was performed in glass-bottomed 35 mm FluoroDishes (WPI) in FluoroBrite DMEM medium (ThermoFisher). DNA was stained with 25 nM SiR-DNA (Spirochrome). Multiple regions of cells were imaged through ten 2.5 μm sections at 5 min intervals for 16 h using the Visitech VT-iSIM super-resolution microscope described above, with a 40x NA1.3 S Fluor Oil objective and Elements 5.21.03 software (Nikon, Japan). A humidified environment at 37°C and 5% CO_2_ was maintained with an Okolab whole microscope and stage-top incubator (Okolab, Italy).

### Image quantification

GFP-TAF5 to SiR-DNA intensity ratios on chromatin in live cell imaging experiments were quantified using ImageJ 2.3.0. Using sum intensity projections, automated Otsu thresholding on the DNA channel defined masks containing the chromatin regions within dividing cells. Water-shedding and manual deletion of chromatin regions from adjacent (non-dividing) cells was then carried out. The intensities of the DNA and GFP signals within the defined regions were then recorded for each time point. The mean cytosolic background signals for SiR-DNA and GFP channels were subtracted from all corresponding data points.

The results were standardised such that the GFP-TAF5/SiR-DNA intensity ratio at the time point immediately preceding nuclear envelope breakdown equaled 1. The first time point at which initial separation of two chromosome masses was evident was taken as anaphase onset. Statistical analysis was carried out in Prism 9.4.1 (GraphPad) using a two-way mixed effects model ANOVA.

### Chromatin isolation and shearing

Cells were fixed in medium with 1% formaldehyde for 10 min at room temperature before quenching with 125 mM glycine for 5 min. Cells were washed once with PBS before adding cold lysis buffer iL1b (from the Auto iDeal ChIP-seq Kit for Transcription Factors, Diagenode C01010058). Asynchronously growing cells (collected by scraping) and mitotic-enriched cells (as described earlier) were suspended at approximately 50 × 10^6^ in 50 ml in buffer iL1b and incubated at 4°C on a roller for 20 min, pelleted, and resuspended in 30 ml buffer iL2 for 10 minutes at 4°C. pelleted material was then resuspended in buffer iS1b with protease inhibitors (100 μl per 3 million cells) and incubated on ice for 10 min. The sample was divided into 270 μl aliquots in Pico sonication tubes (Diagenode C30010016) and sheared using a Bioruptor Pico (Diagenode). Asynchronous samples were sheared for 8 cycles of 15 s with 30 s breaks, and mitotic samples for 5 cycles of 30 s, to yield DNA fragments of 100 - 500 bp. Samples were then centrifuged at 16,000 g for 10 min and the supernatant collected and pooled. If phosphatase inhibitors were used, PhosSTOP (Roche) was added to buffers iL1b, iL2 and iS1b. Sheared chromatin in aliquots of 210 μl (at approximately 1.5 mg protein/ml based on NanoDrop (ThermoFisher) quantification at A_280_) was snap frozen in liquid nitrogen and stored at -80°C.

### SDS-PAGE and immunoblotting

Sheared chromatin was diluted in NuPAGE LDS sample buffer (ThermoFisher) with 10% DTT and 1% Benzonase and incubated at 95°C for 15 min. For analysis of precipitated proteins in CIDOP experiments, elution from washed MagneGST beads was carried out by boiling in in NuPAGE LDS buffer for 10 min.

Samples were electrophoresed on 4 - 12% Bis-Tris gels and analyzed by immunoblotting using standard procedures.

### ChIP-seq

The Auto iDeal ChIP-seq for Transcription Factors protocol (Diagenode) was followed using 200 μl sheared chromatin and the IP-Star Compact Automated System and the direct ChIP iPure 200 method (Diagenode). The antibody coating step was for 3 h, immunoprecipitation for 15 h, and washes for 5 min each at 4°C. Antibodies were used at 3 - 8 μg and DiaMag protein A-coated magnetic beads between 10 and 25 μl accordingly. As crosslinks were reversed, Proteinase K was also added. After crosslink reversal, RNase A was added for 30 min at 37°C. DNA was purified from the ChIP samples (and from an aliquot of sheared chromatin as “input”) using iPure beads (Diagenode) as stated in the Auto iDeal protocol.

Finally, samples were quantified using a Qubit 4 fluorometer and the dsDNA HS Assay kit (ThermoFisher). ChIP libraries were prepared using NEBNext Ultra II DNA Library Prep Kit for Illumina per manufacturer’s instructions (NEB). Sequencing was carried out on Illumina’s NextSeq 500 platform to produce 75 bp single-end reads at a minimum of 50 million reads per sample. ChIP-seq was carried out in 2 (for input and H3K4me2/3) or 3 (for input and H3T3ph) biological replicates.

### CIDOP-seq

Sheared chromatin (100 μl aliquots) was supplemented with 95 μl 0.1% BSA, 50 mM Tris, 434 mM NaCl, 0.5% NP-40, 2 mM DTT, 1 mM ZnCl_2_, pH 7.4, with protease inhibitors (Diagenode). Chromatin was then precleared with MagneGST magnetic beads (Promega) for 1 h at 4°C, and then incubated with approximately 75 μg wild type or M880A recombinant GST-TAF3-PHD overnight at 4°C. MagneGST beads were washed with PB300 (50 mM Tris pH 7.4, 300 mM NaCl, 0.5% NP-40, 2 mM DTT) then added at 12.5 μl per sample for 1 h. The beads were then washed three times for 10 min each at 4°C in buffer PB300, once in 10 mM Tris, pH 7.4, then resuspended in 100 μl buffer iE1 (Diagenode) and incubated rotating for 30 min at room temperature before removing the beads. For the input, 2 μl of precleared chromatin was added to 98 μl buffer iE1 (Diagenode). To samples and inputs, 4 μl iE2 was added then Proteinase K was added for 4 hours at 65°C before adding RNase A for 30 min at 37°C. DNA was purified as for ChIP-seq samples following the Diagenode iPure protocol using reagents from the Auto iDeal ChIP-seq kit (Diagenode). DNA quantification, library construction and sequencing were performed as for ChIP-seq. Two (asynchronous) or three (mitotic) biological replicates were produced for each condition (input, GST-TAF3-PHD WT, GST-TAF3-PHD M880A).

### Sequencing data formatting and visualisation

FastQC v0.11.7 (http://www.bioinformatics.babraham.ac.uk/projects/fastqc/) and MultiQC v1.7216 were used to assess the quality of sequencing reads. Reads with a minimum Phred score of 20 were aligned to reference genome GRCh38.p12 (GCA_000001405.27) using Bowtie2 v2.3.4.2 (Langmead and Salzberg, 2012) with the --very-sensitive option. Samtools v1.9 (Li et al., 2009) was used to create a bam file of the alignments in addition to sorting and indexing. Bam files were converted to bigwigs using bedtools v2.29.2 genomeCoverageBed (Quinlan and Hall, 2010) followed by bedSort and bedGraphToBigwig (UCSC Genome Browser ‘kent’ bioinformatic utilities; http://genome.ucsc.edu/) for data visualisation in IGV v2.4.16 (Robinson et al., 2011). Ideograms were created using the R package KaryoploteR (Gel and Serra, 2017) and the density of ChIP-seq reads in bedgraph format were plotted. ChIP-seq replicates were merged using Sambamba v0.7.1 (Tarasov et al., 2015) and normalised to the appropriate inputs using deepTools2 bamCompare (Ramirez et al., 2016), scaling by read depth and reporting the output as a ratio in bigwig format.

Peak calling of ChIP-seq data was carried out using MACS2 v2.1.1.20160309 (Zhang et al., 2008) with the --broad option and standard parameters. Peaks called in more than one replicate were retained using bedtools intersect, sort and merge tools plus bedops v2.4.39 (Neph et al., 2012) using option -- everything.

### Sequencing data analysis

Metagene plots and heatmaps were plotted from input-normalised bigwigs using deepTools2 computeMatrix with the reference-point option, followed by either plotProfile or plotHeatmap with standard parameters. TSSs were from the UCSC Table Browser and H3K4me2/3 peaks called as described above.

To define the H3T3ph enriched centromere proximal regions, we used a binarisation approach because standard peak calling algorithms perform poorly on broad domains such as centromeric H3T3ph. Bam files were binarized using ChromHMM BinarizeBam with the -center option (Ernst and Kellis, 2012) with a bin size of 200 bp to determine H3T3ph signal per bin. The input for each sample was used as the control data set. Binarised H3T3ph files of all three replicates were then merged and analysed. The largest contiguous block of H3T3ph positive bins on each chromosome was identified (which corresponded to the centromere in each case). Then, to define the two ends of the entire H3T3ph domain on each chromosome, we extended outwards from the contiguous block. The extension continued until 3000 consecutive bins (600 kb) showing no H3T3ph signal were reached. Using this method, 95.2% of the H3T3ph positive bins across all chromosomes were included in the centromere proximal regions. The TSSs within this region were then identified and used in the analysis of centromere-proximal regions in CIDOP-seq data. To complement this analysis, we performed the same procedure for TSSs outside this region (i.e. chromosome arms).

The Pearson correlation of CIDOP-seq or ChIP-seq samples was calculated using deepTools2 multiBigwigSummary bins and input-normalised bigwigs, with standard parameters, followed by plotCorrelation --corMethod pearson, with standard parameters, to produce a heatmap.

### Statistical Analyses

Statistical analyses for binding and kinase assays were carried out in Prism 9.4.1 (GraphPad) with non-normalised data using a two-way repeated measures ANOVA or mixed effects model, followed by Dunnett’s multiple comparisons test. Adjusted p values are reported.

### Additional Files

**Video 1, related to Figure 4**. Live imaging of a GFP-TAF5-expressing wild type HeLa cell progressing through mitosis.

**Video 2, related to Figure 4**. Live imaging of a GFP-TAF5-expressing Haspin-knockout HeLa cell progressing through mitosis.

For both videos:

Left panel: grayscale image of GFP fluorescence. Middle panel: grayscale image of DNA stained with SiR-DNA. Right panel: merged images of GFP (green) and DNA (magenta). Images were taken every 5 min.

