## Supplemental Figures and Table for "Release of Histone H3K4-reading transcription factors from chromosomes in mitosis is independent of adjacent H3 phosphorylation"

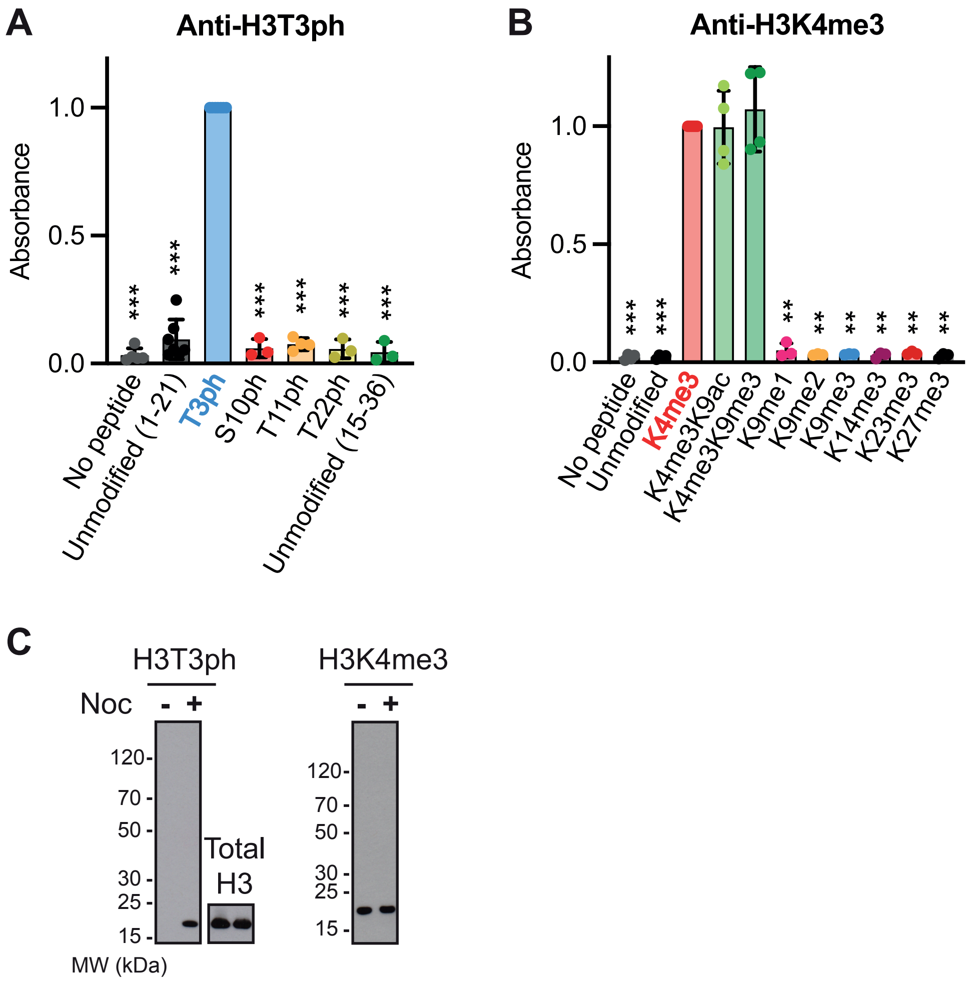


**Figure S1, related to Figure 1. Additional antibody characterisation**

**A**. H3T3ph antibody B8634 binding to H3 peptides with various phosphorylations detected by ELISA (n = 3 to 7).

**B**. H3K4me3 antibody C42D8 binding to H3 peptides with various methylations detected by ELISA (n = 3 to 6).

Data were normalized to the mean signal of the antibodies on the respective target peptide. Bars represent the mean ± SD. Statistical analysis was carried out using non-normalised data, *** $p < 0.0001$, ** $p < 0.001$, * $p < 0.01$, when compared to binding to the expected modification (H3T3ph or H3K4me3).

**C**. Immunoblotting of asynchronous and nocodazole-treated (mitotic) HeLa whole cell lysates with H3T3ph antibody B8634 and H3K4me3 antibody C42D8. The total Histone H3 loading control was carried out in parallel with anti-H3T3ph in the same experiment.

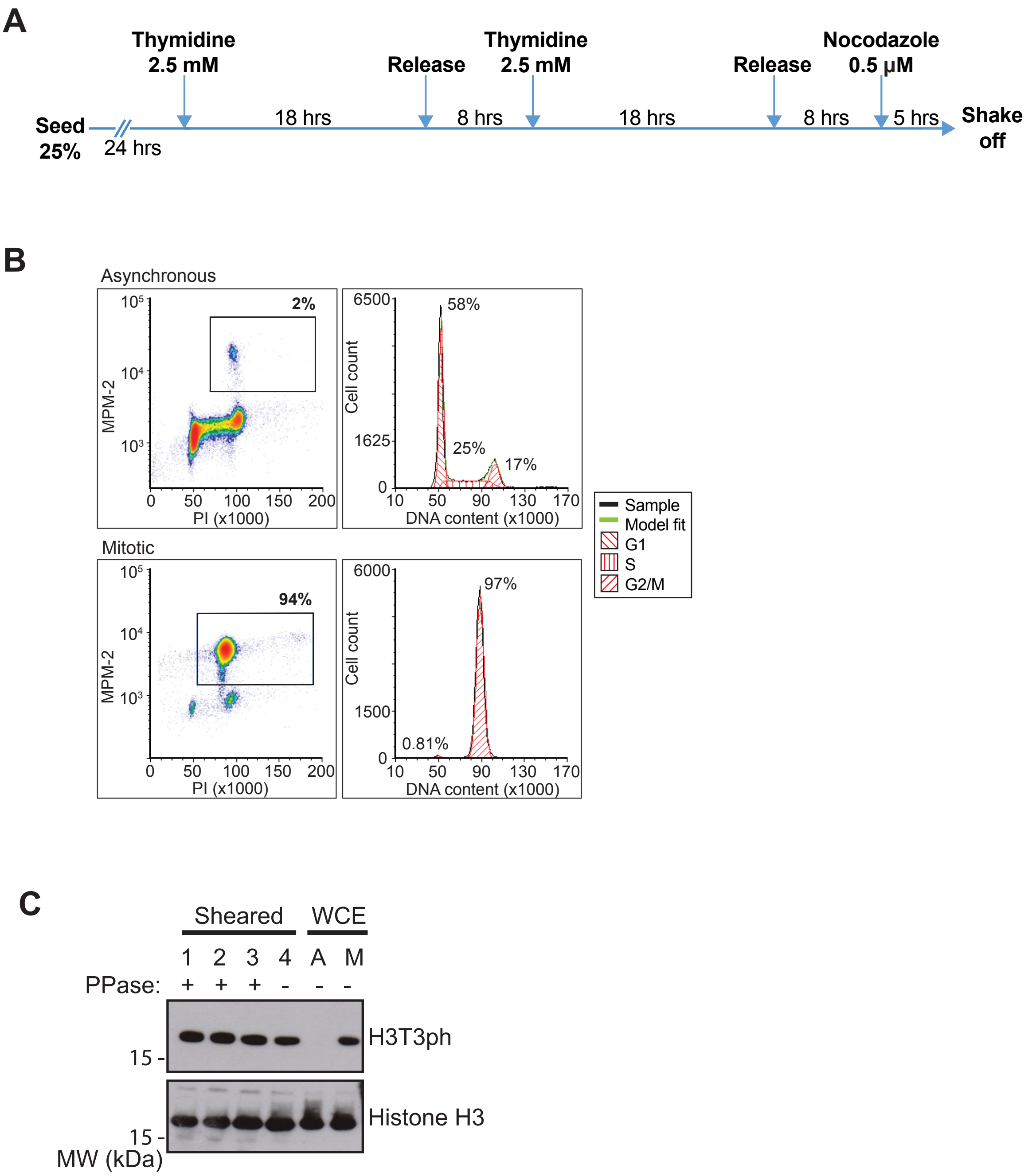


**Figure S2, related to Figure 2. Mitotic enrichment of HeLa cells for ChIP-seq**

**A**. Schematic of synchronization method.

**B**. Representative flow cytometry results of asynchronous cells (top) or cells after synchronization in mitosis (bottom). Percentages indicate the proportion of cells in different cell cycle stages. The mitotic populations, as defined by MPM-2 and propidium iodide (PI) staining, are boxed.

**C**. Immunoblot of sheared chromatin (preparations 1 to 4) or whole cell extract (WCE) from asynchronous (A) or synchronized mitotic (M) HeLa cells probed for H3T3ph or total H3. Whether protein phosphatase inhibitors were included during chromatin preparation is indicated.

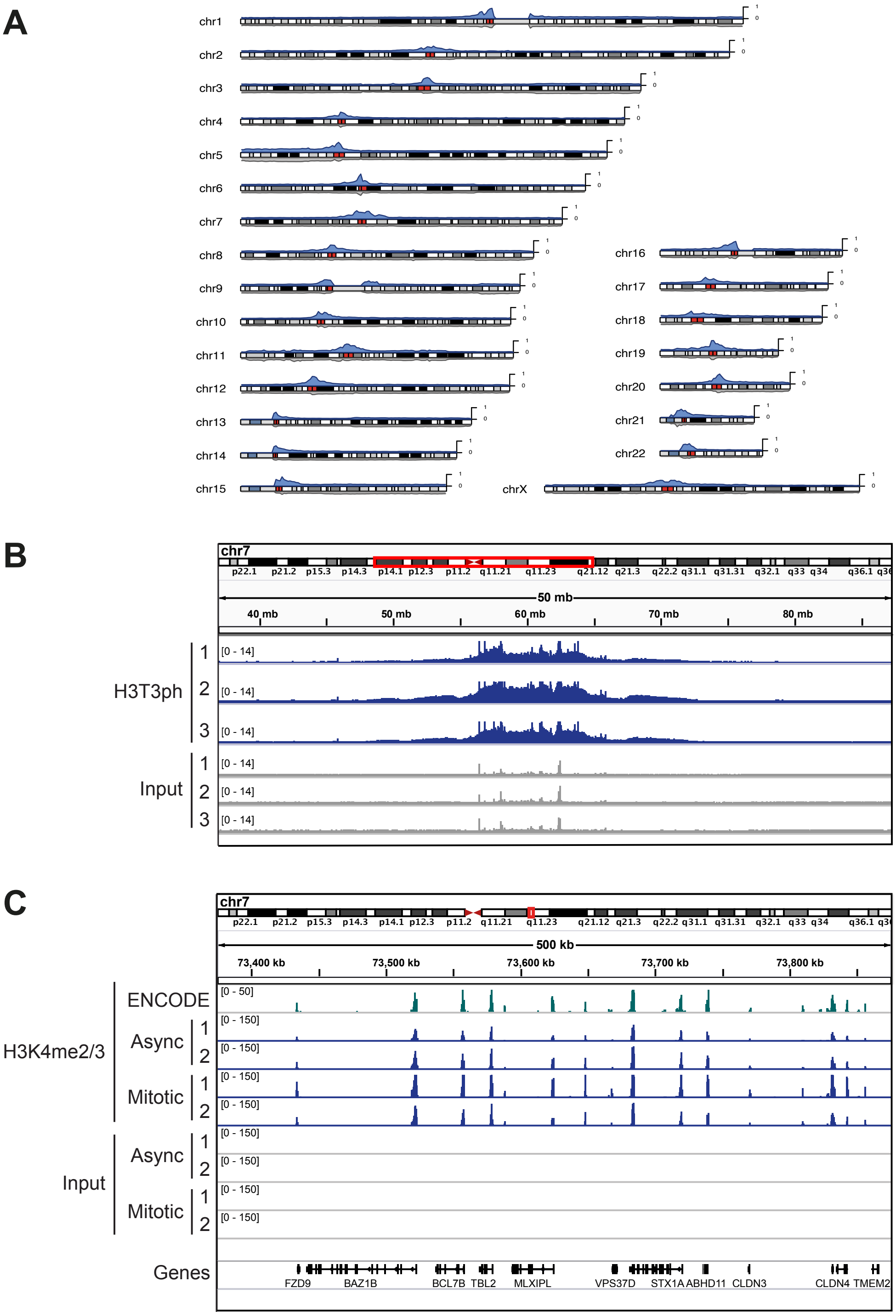


**Figure S3, related to Figure 2. ChIP-seq results and comparison with previous ENCODE data**

**A.** Ideograms showing enrichment of H3T3ph at the centromeres of all human chromosomes. Blue tracks represent H3T3ph read data from a single replicate aligned to GRCh38.p12. Corresponding input reads are plotted in gray. Gaps at the centromeres of chromosomes 1 and 9 reflect incomplete sequence information for these regions in GRCh38.p12.

**B.** Integrative Genomics Viewer (IGV) tracks of H3T3ph ChIP-seq for a 50 MB region of chromosome 7 encompassing the centromere (GRCh38.p12). Read coverages for replicates 1 to 3 are shown (H3T3ph in blue, inputs in grey). Reads mapping to multiple sites were randomly assigned.

**C.** IGV ChIP-seq tracks for a 500 kb region of chromosome 7. H3K4me3 ChIP-seq from the ENCODE project (ENCFF489CIY; fold change over control; green), H3K4me2/3 ChIP-seq replicates 1 and 2 generated in the present study (blue), and the corresponding inputs (gray), for both the asynchronous and mitotic HeLa cells are shown by read coverage. The bottom row shows genes present in this region.

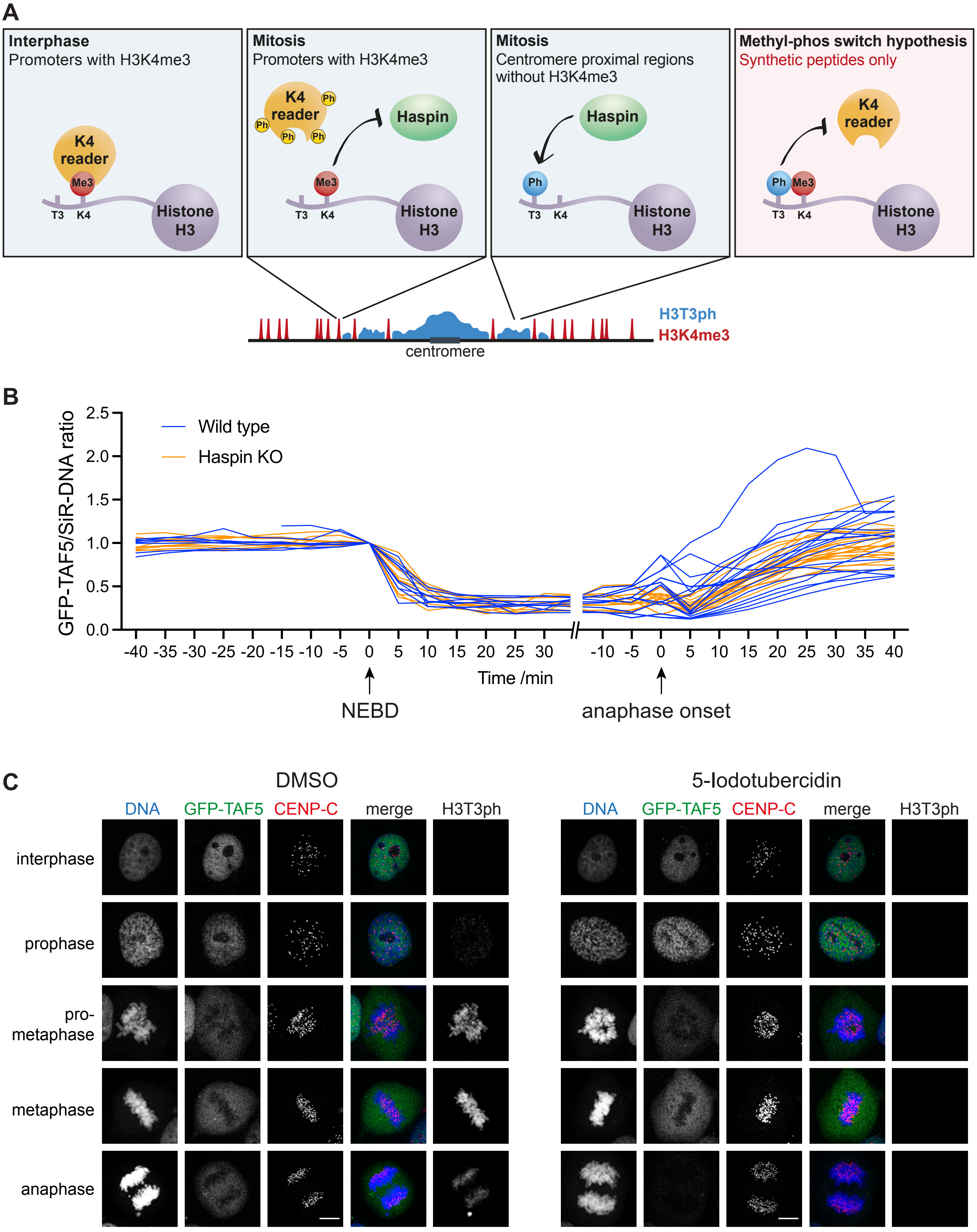


**Figure S4, related to Figure 4. Haspin knock out or inhibition does not influence the displacement of GFP-TAF5 from chromosomes in mitosis**

**A**. Schematic showing the effects of H3T3ph on H3K4me3-reader binding.

**B**. Quantification of GFP-TAF5/SiR-DNA ratio during live imaging of GFP-TAF5-expressing wild type and Haspin knockout HeLa cells. DNA was stained with SiR-DNA, images were taken every 5 min, and times are stated in minutes before and after nuclear envelope breakdown (NEBD) and anaphase onset as appropriate. Traces for individual wild type (n = 9) and Haspin knockout (n = 10) HeLa cells imaged in 3 separate experiments are shown.

**C**. Immunofluorescence microscopy (with formaldehyde fixation) for DNA (blue), GFP-TAF5 (green), CENP-C (centromeres, red), and H3T3ph (gray) in untreated and 5-iodotubercidin (Haspin inhibitor) treated U2OS cells. Scale bars = 10 µm.

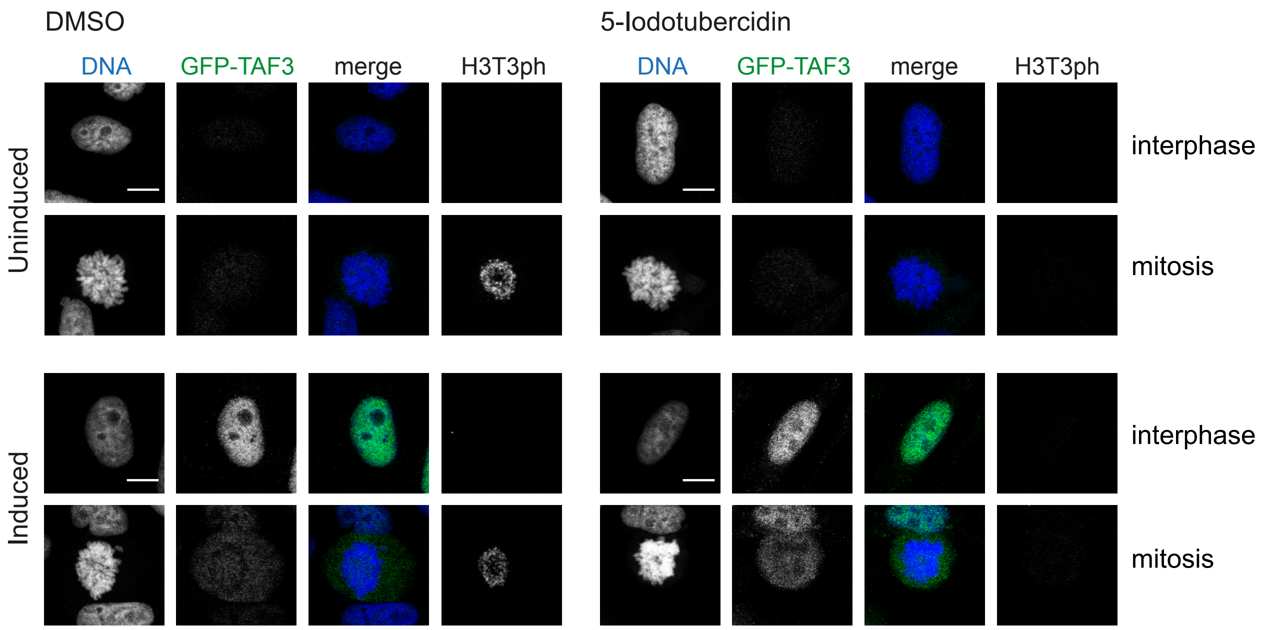


**Figure S5, related to Figure 5. Haspin inhibition does not influence the displacement of GFP-TAF3 from chromosomes in mitosis**
Immunofluorescence microscopy (with formaldehyde fixation) for DNA (blue), GFP (green), and H3T3ph (gray) in HeLa cells inducibly expressing GFP-TAF3 that were treated, or not treated, with the Haspin inhibitor 5-iodotubercidin. GFP-TAF3 expression was induced with doxycycline where indicated. Scale bars = 10 µm.

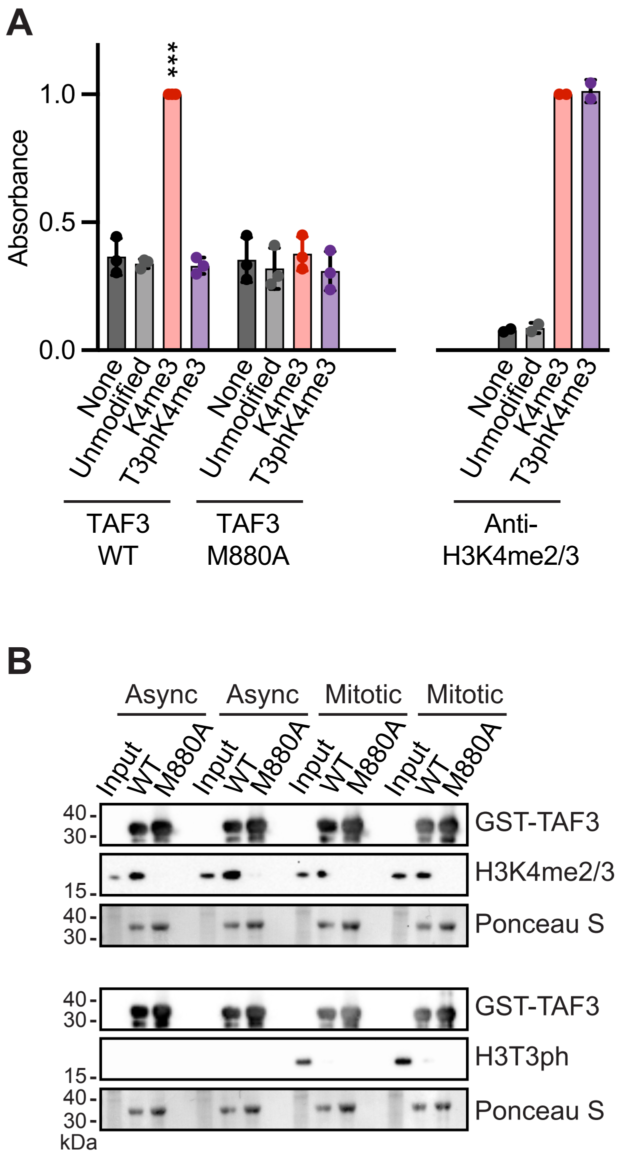


**Figure S6, related to Figure 6. The influence of H3T3ph on GST-TAF3 PHD finger binding to H3K4me3**

**A**. Left: wild type (WT) and M880A mutant GST-TAF3 PHD finger binding to H3 peptides with various modifications (n = 3). Right: Controls showing H3K4me2/3 antibody C42D8 binding to the same H3 peptides (n = 2). Data were normalized to the mean signal of GST-TAF3 or H3K4me2/3 antibody binding to H3K4me3 peptide. Bars represent mean ± SD. Statistical analysis was carried out where n > 2, using non-normalised data. *** $p < 0.0001$, ** $p < 0.001$, * $p < 0.01$, when compared to binding in the absence of peptide.

**B**. Wild type, but not M880A mutant, GST-TAF3 PHD finger immunoprecipitates Histone H3 carrying H3K4me2/3, but not H3T3ph, from asynchronous and mitotic HeLa cell extracts (n = 2 independent samples).

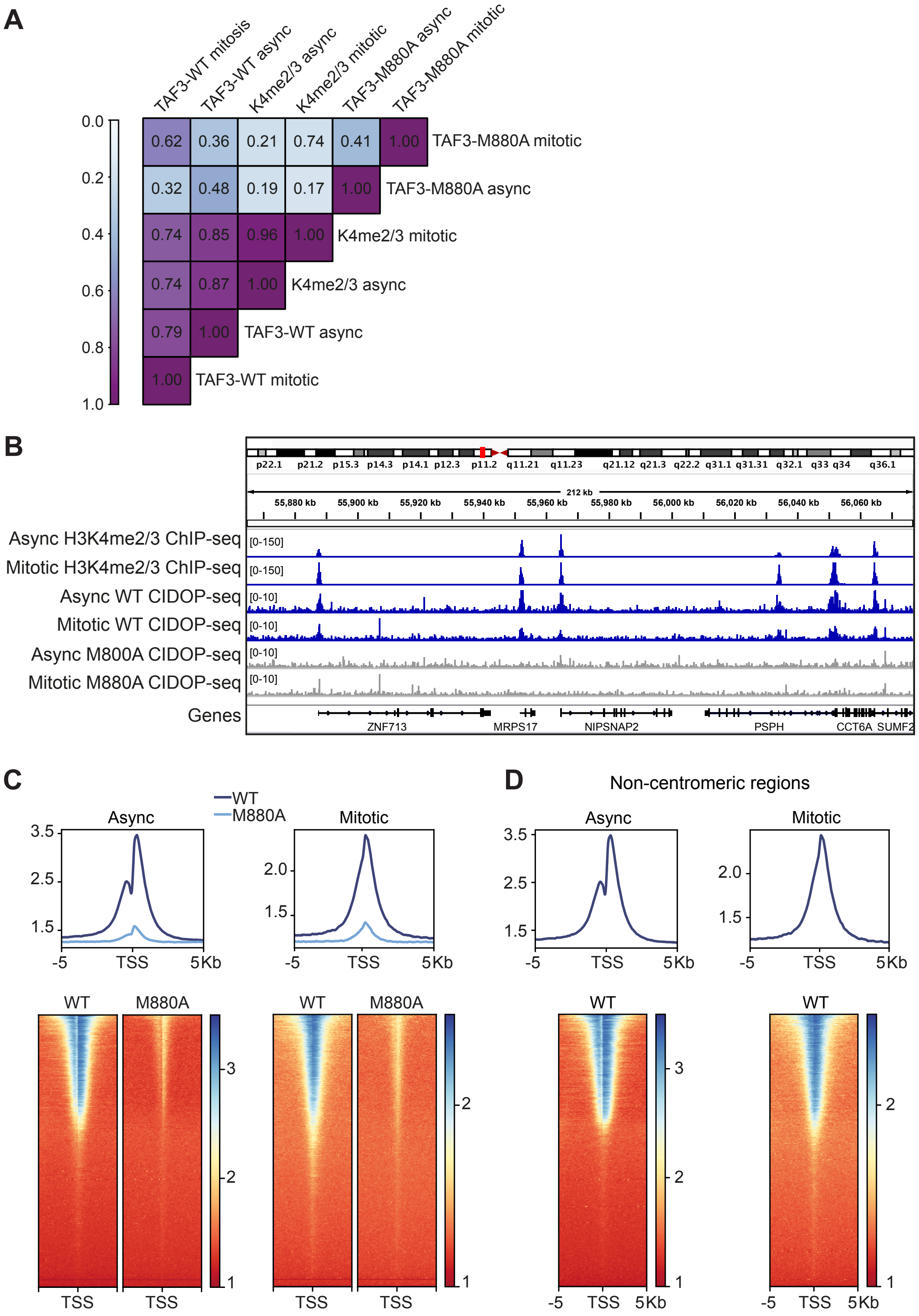


**Figure S7, related to Figure 6. CIDOP-seq of GST-TAF3 PHD finger binding to chromatin from asynchronous and mitotic HeLa cells**

**A**. Pearson correlation coefficients between genome wide read coverage of TAF3 PHD CIDOP-seq (WT or M880A mutant) and H3K4me2/3 ChIP-seq for both asynchronous and mitotic-enriched HeLa cells.

**B**. Representative IGV tracks of a 212 kb region of chromosome 7, showing TAF3 PHD (WT or M880A mutant) CIDOP-seq and H3K4me2/3 ChIP-seq for asynchronous and mitotic-enriched HeLa cells.

**C**. WT and M880A mutant TAF3 PHD CIDOP-seq enrichment at TSSs genome wide. Results from both asynchronous and mitotic-enriched HeLa cells are shown metagene plots (top) and as heatmaps (bottom). Regions of 10 kb centered at TSSs are shown.

**D**. TAF3 PHD enrichment across TSSs that are not centromere-proximal (i.e. those on chromosome arms where H3T3ph is low). Metagene plots (top) and heatmaps (bottom) show TAF3 PHD binding at TSSs for both asynchronous and mitotic-enriched CIDOP-seq HeLa cells.

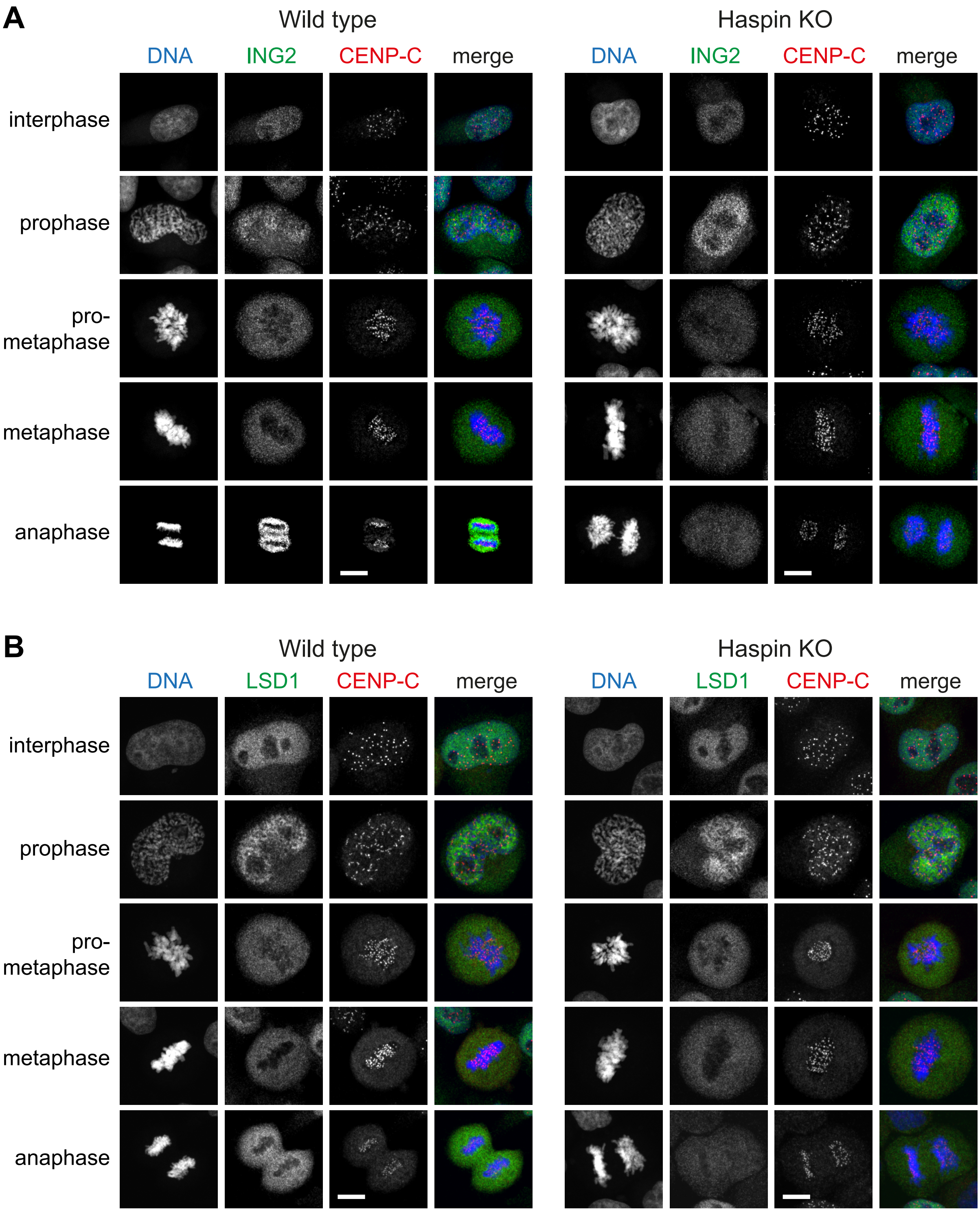


**Figure S8, related to Figure 7. Haspin knockout does not influence the displacement of endogenous ING2 or LSD1 from chromosomes in mitosis**

**A**. Immunofluorescence microscopy (with formaldehyde fixation) for DNA (blue), ING2 (green), CENP-C (centromeres, red), and H3T3ph (gray) in wild type and Haspin knockout HeLa cells.

**B**. As for A, but staining for LSD1 (green). Scale bars = 10 µm.

**Supplemental Table 1**
Mammalian H3K4-reading proteins that are displaced by H3T3ph *in vitro*.

| Reader protein | Histone mark | Reference |
|---|---|---|
| **AIRE** PHD1 domain | H3K4me0/1 | (Chignola et al., 2009; Garske et al., 2010) |
| **BHC80** (**PHF21A**) PHD domain | H3K4me0 | (Garske et al., 2010; Varier et al., 2010) |
| **BHC80L** (**PHF21B**) PHD domain | H3K4me1 | (Basu et al., 2020) |
| **BPTF** PHD domain | H3K4me3 | (Fuchs et al., 2011; Varier et al., 2010) |
| **BRPF2** (**BRD1**) PHD1 domain | H3K4me0 | (Qin et al., 2011) |
| **CFP1** PHD domain (**Spp1** in *S. pombe*) | H3K4me2/3 | (He et al., 2019) |
| **CHD1** DCD domain | H3K4me3 | (Flanagan et al., 2005; Fuchs et al., 2011; Gatchalian et al., 2016) |
| **CHD4** PHD domain | H3K4me0 | (Jain et al., 2020; Mansfield et al., 2011) |
| **CHD5** PHD domain | H3K4me0 | (Jain et al., 2020; Oliver et al., 2012) |
| **DIDO** PHD domain (**Bye1** in *S. cerevisiae*) | H3K4me3 | (Gatchalian et al., 2013; Gatchalian et al., 2016; Jain et al., 2020; Kinkelin et al., 2013; Tencer et al., 2017) |
| **DNMT3A** ADD domain | H3K4me0/1 | (Noh et al., 2015; Zhang et al., 2010) |
| **DNMT3B** ADD domain | H3K4me0/1 | (Noh et al., 2015; Zhang et al., 2010) |
| **DNMT3L** ADD domain | H3K4me0 | (Noh et al., 2015) |
| **DPF2** PHD domain | H3K4me0 | (Jain et al., 2020) |
| **ING1** PHD domain | H3K4me3 | (Gatchalian et al., 2016) |
| **ING2** PHD domain | H3K4me3 | (Garske et al., 2010; Varier et al., 2010) |
| **ING4** PHD domain | H3K4me3 | (Varier et al., 2010) |
| **KDM4A** DTD domain | H3K4me3 | (Gatchalian et al., 2016; Su et al., 2016) |
| **KDM5B** PHD1 domain | H3K4me0 | (Klein et al., 2014) |
| **KDM5B** PHD3 domain | H3K4me3 | (Klein et al., 2014) |
| **KDM7A** PHD domain | H3K4me3 | (Jain et al., 2020) |
| **MLL1** SET domain | H3K4me0 | (Southall et al., 2009) |
| **MLL5** PHD domain | H3K4me3 | (Ali et al., 2013; Jain et al., 2020) |
| **ORC1b** PHD domain (*Arabidopsis*) | H3K4me0 | (Li et al., 2016) |
| **PHF8** PHD domain | H3K4me3 | (Gatchalian et al., 2016) |
| **PHRF1** PHD domain | H3K4me0 | (Jain et al., 2020) |
| **RAG2** PHD domain | H3K4me3 | (Fuchs et al., 2011; Garske et al., 2010; Gatchalian et al., 2016) |
| **SGF29** tandem Tudor domain | H3K4me3 | (Shanle et al., 2017) |
| **SP140** PHD domain | H3K4me0 | (Zhang et al., 2016) |
| **TAF3** PHD domain | H3K4me2/3 | (Gatchalian et al., 2016; Kungulovski et al., 2016; Shanle et al., 2017; Varier et al., 2010) |
| **TRIM66** PHD domain | H3K4me0 | (Jain et al., 2020) |
| **WDR5** WD40 repeats | H3K4me0/1/2/3 | (Couture et al., 2006; Klingberg et al., 2015) |
